# Decoding the transcriptional response to ischemic stroke in young and aged mouse brain

**DOI:** 10.1101/769331

**Authors:** Peter Androvic, Denisa Belov Kirdajova, Jana Tureckova, Daniel Zucha, Eva Rohlova, Pavel Abaffy, Jan Kriska, Miroslava Anderova, Mikael Kubista, Lukas Valihrach

## Abstract

Ischemic stroke is one of the leading causes of mortality and major healthcare and economic burden. It is a well-recognized disease of aging, yet it is unclear how the age-dependent vulnerability occurs and what are the underlying mechanisms. To address these issues, we performed a comprehensive RNA-Seq analysis of aging, ischemic stroke and their interaction using a model of permanent middle cerebral artery occlusion (MCAO) in 3 and 18 month old female mice. We assessed differential gene expression across injury status and age, estimated cell type proportion changes, assayed the results against a range of transcriptional signatures from the literature and performed unsupervised co-expression analysis, identifying modules of genes with varying response to injury. We uncovered selective vulnerability of neuronal populations and increased activation of type-I interferon (IFN-I) signaling and several other inflammatory pathways in aged mice. We extended these findings via targeted expression analysis in tissue as well as acutely purified cellular populations to show differential temporal dynamics of IFN-I signaling between age groups and contribution of individual cell types. Together, these results paint a picture of ischemic stroke as a complex age-related disease and provide insights into interaction of aging and stroke on cellular and molecular level.

**Graphical summary:** 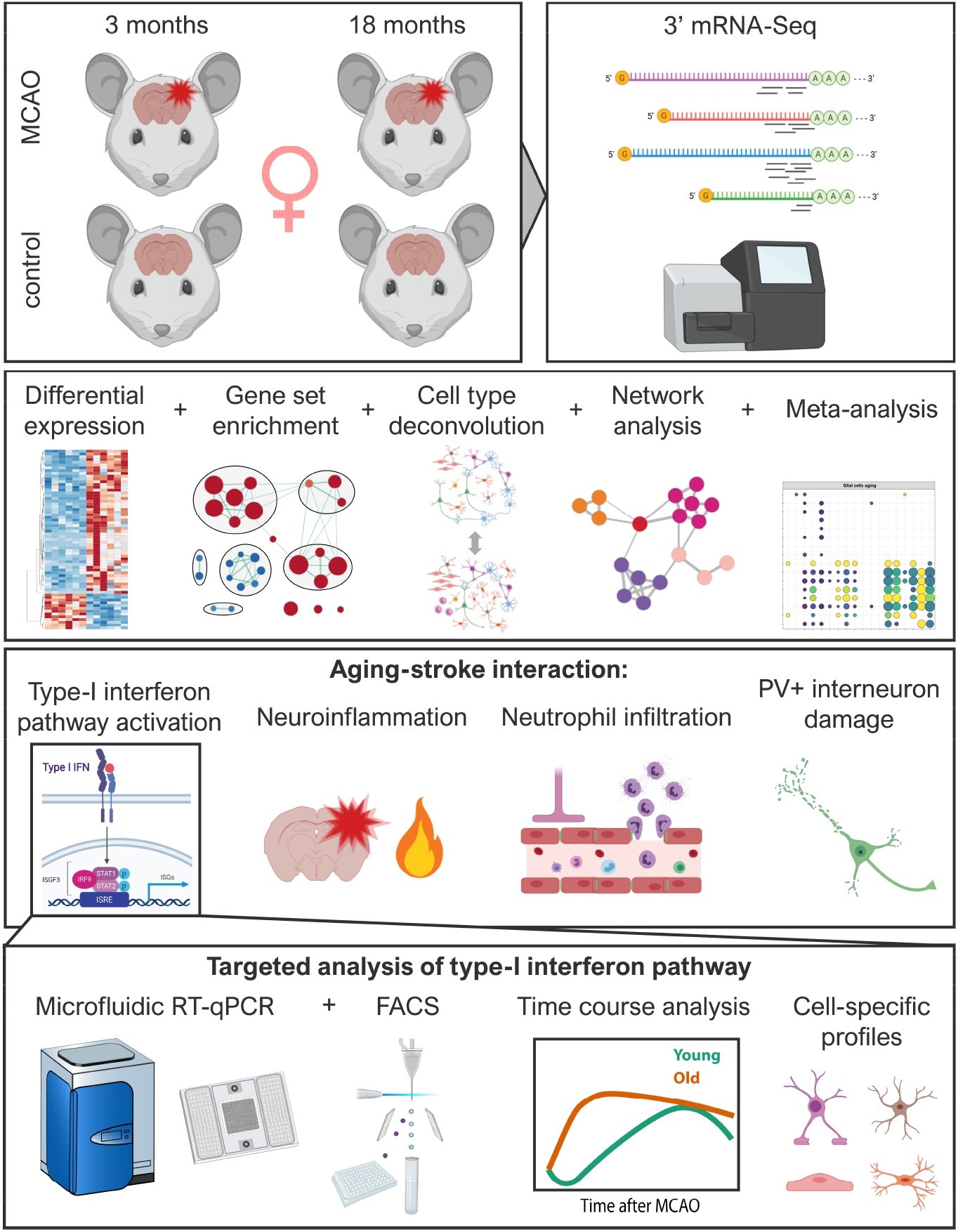

## Introduction

Cerebral stroke affects nearly 17 million people per year worldwide, with a death rate of 5.9 million^1^, making it the second leading cause of death in the developed world^2^. In addition, stroke is the primary cause of long-term disability^3^ and represents a major healthcare and economic burden^4^. Ischemic stroke is caused by a loss of blood flow to the brain and accounts for ~87% of all strokes. The remaining ~13% of stroke cases are hemorrhagic and are caused by blood leakage into the brain^5^.

Pathophysiology of ischemic stroke is complex, and involves various mechanisms including disruption of blood-brain barrier (BBB), excitotoxicity, inflammation, oxidative damage, ionic imbalances, apoptosis, angiogenesis and neuroprotection. The ultimate result of ischemic cascade is neuronal death along with an irreversible loss of neuronal function^6,7^. Two main approaches considered to treat acute ischemic stroke are reperfusion (restoration of blood flow) and neuroprotection (protection of neurons from ischemic injury)^8,9^. Despite substantial efforts invested into research and development of neuroprotective strategies, with more than 1000 drugs investigated and 100 tested in clinical trials^10^, early clot lysis with recombinant tissue plasminogen activator remains the sole approved therapy^11^.

One of the reasons for this translational roadblock is that most preclinical studies have used only young animals, despite tremendous evidence that ischemic stroke is a disease of aging^2,12–14^. 75-89% of strokes occur in people aged > 65 years, and for each decade after the age of 55 years the stroke rate doubles^2^. Along with higher incidence, aged patients have higher mortality, suffer more severe deficits and recover slower than younger patients^15,16^. In addition, sex modifies the influence of age; although ischemic burden is higher in men throughout most of the lifespan, elderly women suffer more strokes and have poorer functional outcomes and quality of life^13,17–20^. These clinical features are largely recapitulated in experimental rodent models of acute ischemic stroke. Old animals of both sexes have higher mortality and more severe neurological deficits than young animals^21–23^. In addition, sex-specific differences are seen throughout the lifespan in infarct volumes, inflammation, blood-brain barrier (BBB) permeability^12,24^ and glial cell reactivity^25,26^.

With rapidly aging human population, the global stroke burden increases. There is a need to understand the age-related mechanisms of ischemic injury to improve our ability to discern which therapeutics can be translated from the bench to the bedside^2^. Previous global gene expression studies of experimental stroke using microarrays^27–34^ and more recently RNA-Seq^35–37^ have provided useful insights into the pathophysiology of ischemic stroke and uncovered many altered molecular pathways^38^. However, few studies included aged animals. One microarray report found that inflammation and synaptic plasticity-related genes had attenuated transcriptional response to stroke in aged mice^39^. Another microarray study identified age-dependent response of DNA-damage and pro-apoptotic genes, but assayed only limited number of genes (442) in pre-selected pathways^40^, while the follow-up study focused solely on angiogenesis^41^. In addition, these studies employed only male animals and clinically less relevant model of transient ischemia^42–46^. The complex factors underlying worsened stroke outcome in the elderly thus remain poorly understood, particularly in females that are often underrepresented in both clinical and pre-clinical stroke research^14,47–49^.

In this study, we aimed to dissect the interaction between stroke and aging at the genome-wide level. We used permanent middle cerebral artery occlusion (MCAO) – a clinically most relevant model of ischemic stroke^42–46^ – on young adult (3 months) and aged (18 months) female mice and analyzed the post-ischemic cortex at 3 days after MCAO using RNA-Seq. We combined differential gene expression and pathway analyses with network and cell type deconvolution approaches and intersected the results with relevant transcriptional signatures from the literature. Our results show age-dependent alterations in processes predominantly associated with inflammation and interferon signaling, as well as the selective vulnerability of specific neuronal subpopulations. We then complemented the results with targeted expression analysis on acutely isolated cell populations and provide detailed insight into temporal dynamics and cell-specific response of interferon signaling pathway in the young and aged post-ischemic brain. Our data provide new insights into the mechanisms of ischemic injury in the aged brain and serve as a publicly available resource for future studies.

## Results

We hypothesized that there are two components leading to a more severe outcome of ischemic stroke in aged animals:

i. changes in the brain environment during normal aging making the aged brain more susceptible to ischemic injury, and
ii. the difference in the response to ischemic challenge between young and aged brain leading to secondary injury and impaired regeneration. To explore both of these components, we performed 3’ mRNA sequencing of parietal cortex isolated from 3 month old and 18 month old female mice (representing young adult and aged animals) at 3 days after MCAO and from their age-matched controls (in total 4 groups, 6 animals per group).

### Aging is accompanied with increased neuroinflammation involving primarily glial cells

Building on our hypotheses, we first explored factors that may contribute to the increased vulnerability of the aged brain to ischemic challenge. We compared differentially expressed (DE) genes between young and aged controls and analyzed them using Gene set enrichment analysis (GSEA)^50^. We found 52 upregulated and only 13 downregulated genes (log_2_FC > 1, p_adj_ < 0.05; Figure S1). GSEA revealed upregulation of defense response-related processes, positive regulation of immune response, and increased secretion of cytokines and protein catabolism (Figure 1A). Concomitantly, downregulated processes mapped to positive regulation of protein polymerization, dendrite development and axon projection, altogether pointing towards increased inflammation and axonal degeneration in the aged brain. To gain further tissue- and cell-specific context of observed transcriptional changes, we searched the literature for transcriptomic datasets related to brain aging, neuroinflammation and stroke, and quantified overlap with our lists of differentially expressed genes (Figures 1B, S2). There was a significant overlap between the genes upregulated within the aged controls and signatures of aged astrocytes (such as *Gfap, Anln, Pcdhb6, C4b, Serpina3n, Lyz2, Neat1, Plin4*)^51,52^, aged microglia (such as *Clec7a, Cst7, Cybb, Lgals3, Mmp12, Spp1, C4b, Ccl8*)^53,54^, aging oligodendrocyte precursor cells (OPCs) (such as *Rab37, Tnfaip2*)^55^, in LPS-treated microglia (such as *Bcl3, C3ar1, Ccl3, Ccl4, Cst7, Cybb, Tnfaip2*)^56,57^ and/or LPS-treated astrocytes (such as *Casp1, Flnc, Mpeg1, Runx1, Serpina3n*)^57^, as well as with genes that are part of the common inflammatory signature (such as *Ptprc, Rab32, Slc11a1, St14, Tep1, Trem2, Tyrobp*)^58^. We also found significant upregulation of genes enriched in bone marrow-derived macrophages (BMDMs) versus brain-resident microglia (such as *Cdkn2a, Itgax, Tep1*), while microglia-enriched genes (such as *Cask, Gda, Nav3, Nrep, Sox4*) were downregulated, indicating the convergence of microglial and macrophagal signatures with aging, as previously suggested^54,59^. Furthermore, aging-upregulated genes strongly overlapped with the neurodegeneration-related transcriptional profile of microglia^59^ and recently identified markers of *Ccl4*-expressing subpopulation of microglia that expand during aging, injury^60^ and neurodegeneration^61^. Overall, these results show that brain aging leads to subtle alterations in the neuroinflammatory environment, involving particularly glial cells. A subset of pro-inflammatory primed microglia and/or astrocytes may confer adverse milieu that may contribute to aggravation of the ischemic injury in aged animals.

**Figure 1.**
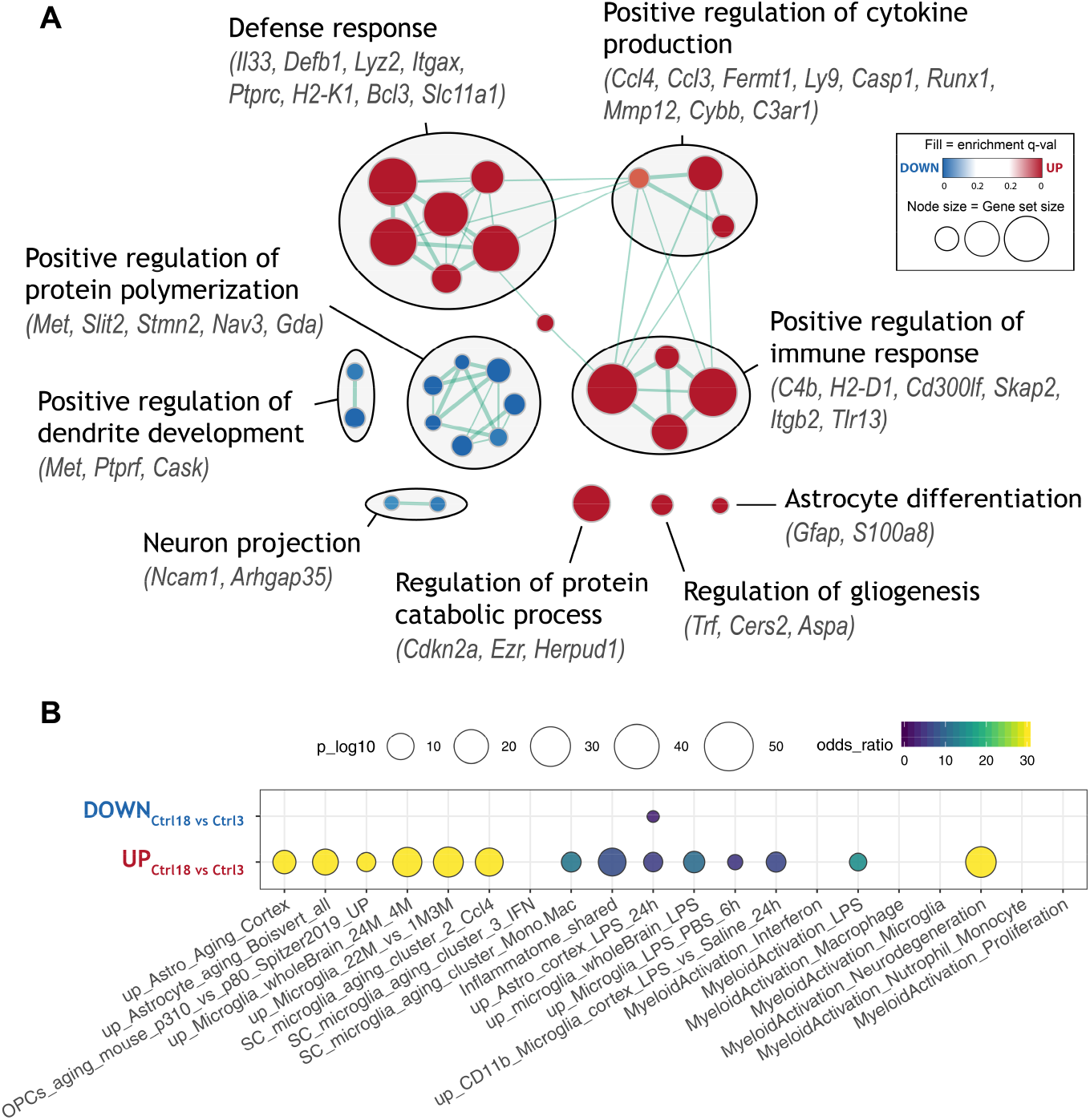
Gene sets with altered expression during normal aging. A) Enrichment map of significantly up- or down-regulated gene ontology (GO) terms between aged (18 months) and young adult (3 months) control mice. Nodes represent gene sets. Highly similar gene sets are connected by edges, grouped in sub-clusters and annotated manually. B) Meta-analysis showing enrichment of selected transcriptional signatures from literature. Several inflammatory and glial cell activation-related states are significantly upregulated. See Table S1 for gene set descriptions.

### Aging alters the magnitude of the transcriptional response to ischemic stroke

Next, we explored transcriptional changes at 3 days after stroke separately in young and aged mice relative to their age-matched controls. A large number of genes were DE in both young (2556) and aged (3435) mice, with a high prevalence of upregulation, both in terms of number of regulated genes and the fold-change (Figures 2A, S3A). There was a substantial overlap between DE genes in young and aged mice, although aged mice differentially regulated more genes, often with a greater magnitude (Figures 2C, S4). Functional analysis with GSEA revealed hundreds of significantly enriched gene ontology (GO) terms. To remove redundancy, we clustered overlapping, functionally related gene sets into the network using Enrichment Map^62^ (Figure S5). This analysis revealed several clusters of stroke-upregulated gene sets related to metabolic activity, reactive oxygen species (ROS) production, extracellular matrix, angiogenesis and two major clusters associated with the cell cycle and immune response (Figure S5). Stroke-downregulated genes were associated with ion channel and synaptic transporter activity, oxidative phosphorylation, neuronal projection and synaptic vesicles production and GO terms “learning and memory” and “locomotory behavior” (Figure S5). Although comparison of the significantly enriched gene sets between young and aged animals revealed substantial overlap, GO terms related to inflammatory response (such as type-I interferon signaling, cytokine production, neutrophil degranulation), cell-cell interactions (such as integrin cell-surface interaction, extracellular matrix organization) and cell-cycle (such as regulation of DNA replication) were upregulated to a greater extent in aged animals after injury (Figures 2B, S3B). Similarly, gene sets associated with synaptic signaling and plasticity, neurotransmitter transport and potassium ion channels were downregulated exclusively, or to a greater extent in aged animals after stroke (Figures 2B, S3B).

**Figure 2.**
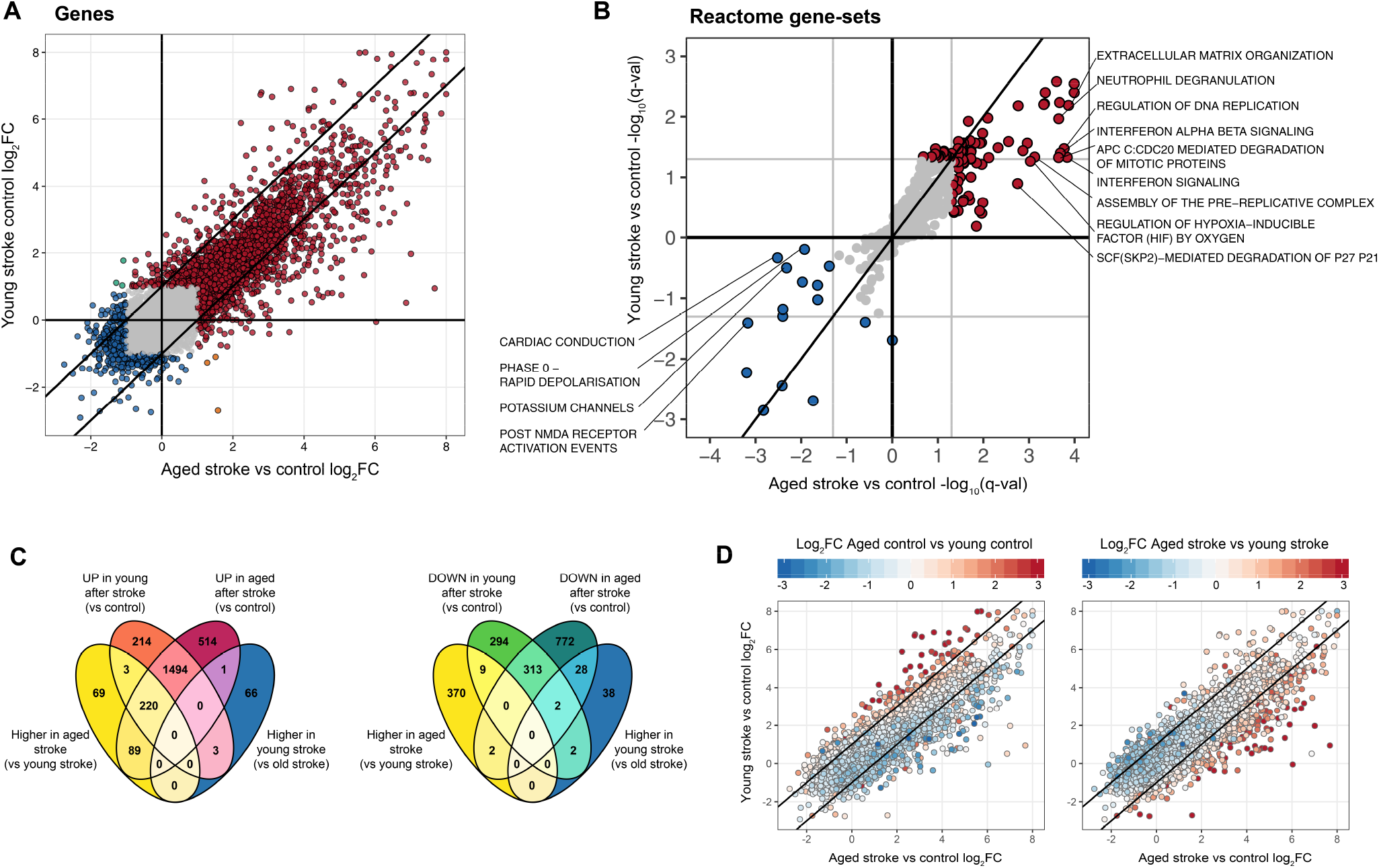
Comparison of differentially expressed genes and gene sets after stroke between young and aged mice. A) Scatter plot comparing stroke-induced log_2_ fold-change in young and aged mice. Genes with |log_2_FC| >1 are highlighted in color. See also Figure S3A. B) Scatter plot comparing stroke-induced alteration of Reactome pathways in young and aged mice. Pathways wih q-val < 0.05 are highlighted in color. Sign depicts UP (+) or DOWN (−) regulation. See also Figure 4S3B and S5. C) Venn diagrams showing overlaps of DE genes between selected pairwise comparisons. See also Figure S4. D) Scatter plots of stroke-induced log_2_ fold-change in young and aged mice. Color maps to log_2_ fold-change between aged and young controls (left) or between aged and young stroke groups (right). Genes with larger stroke-induced upregulation in young mice tend to increase in expression during normal aging.

These results suggest that in addition to common genes, distinct cellular processes are activated and repressed in response to ischemic impact in aged brain. Surprisingly, almost no functional gene sets were induced or repressed exclusively in young animals (Figures 2B, S3B).

### Stroke does not activate exclusive neuroprotective pathways in young compared to aged mice

Since we did not identify any uniquely responsive gene sets in young mice, we took a step back and focused on individual genes that showed a greater stroke-induced upregulation in young mice compared to the aged group (“more up MCAO3”). We postulated that such genes may display a neuroprotective effect (considering the better outcome of stroke in young animals), and serve as possible targets for pharmacological activation in aged patients. We found that although some genes do show greater upregulation upon MCAO in young mice (log_2_FC_young_ ≫ log_2_FC_aged_), almost all of them are also up-regulated in aged controls compared to young controls (Figure 2D, left) and their abundance levels in young stroke group do not rise above the levels in aged stroke group (Figure 2D, right). Considering the inflammatory nature of these genes, the strong overlap with markers of aged and activated microglia^60,61,63^ (Figure S2), as well as the presence of reactive astrocyte marker (*Gfap*), it is apparent that these genes reflect age-induced glial activation, which is partially saturated after stroke in aged mice.

These results suggest that the group of genes showing stronger activation in young animals (after stroke) does not involve exclusive neuroprotective pathways, but rather reflects the resting baseline level of microglia and astrocytes in young control animals and a more polarized baseline state in aged control animals.

### Combination of aging and stroke leads to massive activation of type-I interferon signaling and aggravated inflammatory response

We then explored the genes showing greater upregulation upon ischemic impact in aged mice, which are likely to exert detrimental effects (“more up MCAO18”). There were more than 400 such genes, of which a large proportion were immune-related (Figure 3A). GO and pathway enrichment analysis revealed T-cell activation, cell adhesion, chemotaxis and leukocyte migration, as the ones of the most strongly enriched functional clusters, suggesting an increased infiltration and activation of peripheral immune cells (Figure 3A). We also found strong enrichment of genes associated with antigen processing and presentation, MHC class I, and cytokine secretion. Several signaling pathways, namely ERK/MAPK signaling and cAMP/cGMP mediated signaling were also significantly enriched, as were the GO terms associated with lipid metabolism, transport of fatty acids and oxidative phosphorylation. Pathway enrichment revealed clusters of extracellular matrix organization and cluster of pathways mediating regulation of cell cycle, suggesting that cellular proliferation may be increased in aged post-stroke mice (Figure 3A). Genes that are more induced by stroke in aged mice also significantly overlapped with several inflammation and aging associated signatures from the literature (Figure S2). This was expected, considering the highly inflammatory nature of the genes in the “more up MCAO18” gene set. However, an interesting feature was the strong overlap with the LPS-induced / A1 pro-inflammatory astrocytic profile (Figure S2). Previously, it has been reported that MCAO induces a beneficial A2 astrocytic profile^64^. Our result suggests this may not be the case in the aged brain, which would be consistent with reports that aging promotes inflammatory A1 profile of astrocytes^51,52^ and accelerates injury-induced astrocyte reactivity^65–67^. Another striking feature was particularly strong overrepresentation of interferon-stimulated genes, indicating that specifically type-I interferon signaling pathway is much strongly activated by stroke in aged animals (Figures 2A, S2).

**Figure 3.**
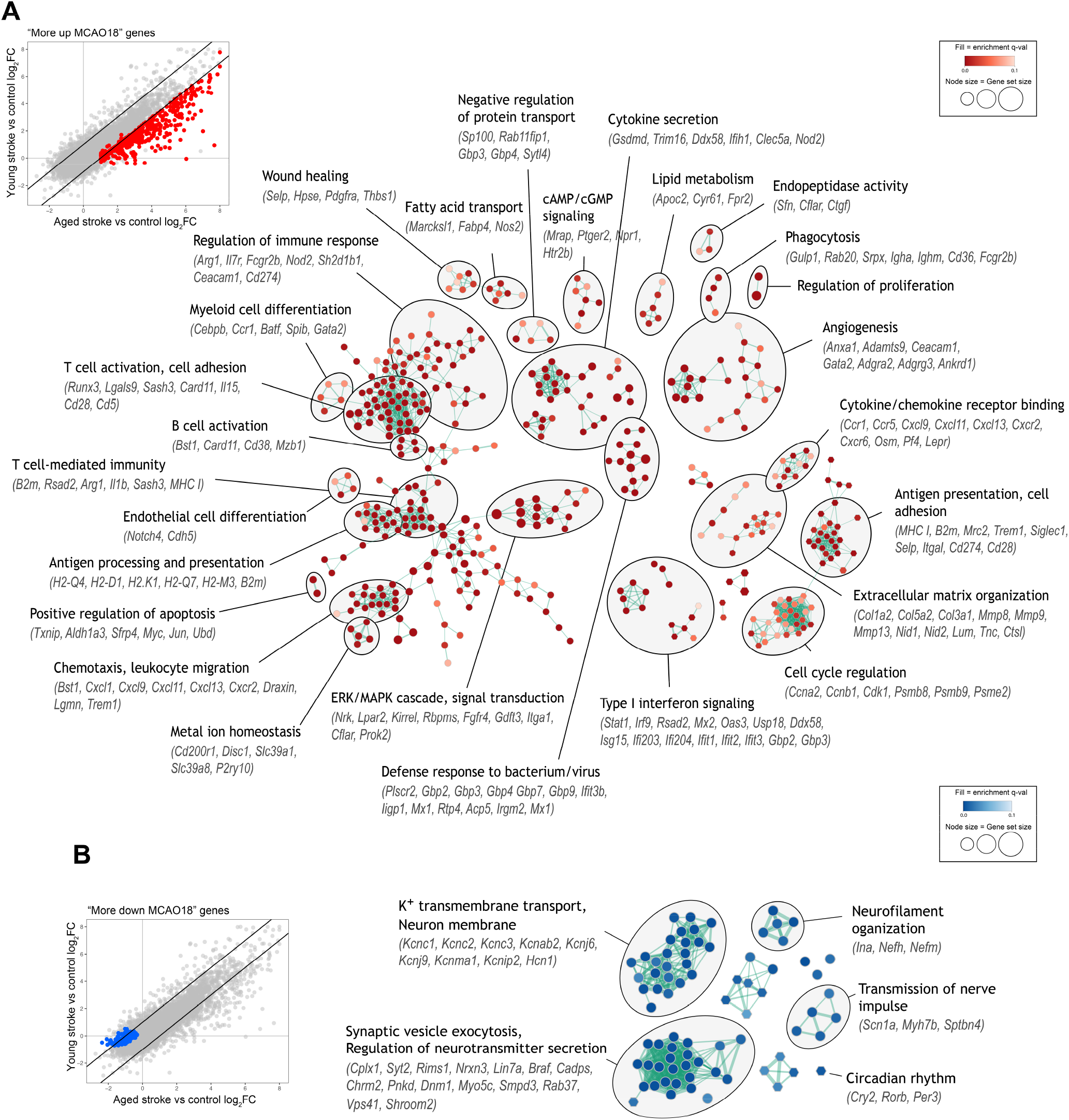
Functional annotation of genes with different quantitative response to stroke between young and aged mice. A) Enrichment map showing significantly enriched GO terms (circles) and pathways (hexagons) for genes with greater upregulation after stroke in aged mice (“more up MCAO18”). Similar gene sets are grouped into sub-clusters and annotated manually. See also Figure S2 for enrichment of gene sets from the literature. B) Same as (A), but for genes with greater downregulation after stroke in aged mice (“more down MCAO18”).

In parallel to larger upregulation, aged mice also downregulated larger number of genes after stroke, often with a greater magnitude (“more down MCAO18”) (Figure 3B). Functional annotation of the “more down MCAO18” gene set revealed significant enrichment of K+ transmembrane transport (*Atp1b1, Hcn1*), voltage-gated K+ channels (*Kcnc1, Kcnc2, Kcnc3, Kcnj6, Kcnj9, Kcnma1, Kcnt1*) and their regulatory subunits (*Kcnab2, Kcnab3, Kcnjp2*), neurofilament proteins (*Nefh, Nefm, Ina*) and genes involved in synaptic vesicle exocytosis regulation and neurotransmitter release (such as *Cadps, Pnkd, Lin7a, Braf, Dnm1, Rims1, Cplx1, Syt2, Nrxn3*) (Figure 3B). These genes mainly localize along the presynaptic and neuron projection membranes, indicating greater axonal damage and impaired synaptic communication in aged post-stroke mice. “More down MCAO18” genes were also enriched with genes involved in the regulation of circadian rhythm (such as *Cry2, Rorb, Per3*). An impact of ischemic stroke on circadian rhythm has been observed before^68^ and our results suggest that aged animals may be more susceptible to its destabilization, which is linked to sleep, mood and post-stroke depression, and may therefore impact recovery^69,70^. Overall, analysis of age-stroke interacting genes revealed an increased neuroinflammatory environment in aged animals, which is connected to higher infiltration and activation of peripheral immune cells, pro-inflammatory cytokine secretion and activation of signaling pathways (ERk/MAPK, type-I interferon) that may contribute to secondary injury. On the other hand, K+ transmembrane channels, neurofilament and synaptic communication proteins were specifically repressed in aged animals, likely reflecting increased axonal damage.

### Transcriptome deconvolution reveals cell type composition changes during aging and after stroke

In order to provide cell-specific context to the observed transcriptional profiles, we assessed relative changes of cell type proportions by computational deconvolution. First, we built a reference of stable cell-specific genes for major CNS cell types (for details see Methods). We then used the *marker-GeneProfile* R package^71^, which summarizes expression of multiple cell-specific genes into a single marker gene profile (MGP), serving as a surrogate for the cell type proportions. To validate the estimates, we used transcriptome deconvolution algorithm CIBERSORT^72^ (see Methods). Except for astrocytes, concordance analysis confirmed the robustness of the results (Figure S6). Closer inspection showed that astrocyte-specific genes do not significantly overlap with any DE geneset (Figure 4B), and cluster into different co-expression modules (see below), which may be due to heterogeneity of astrocytic response or intrinsic transcriptional regulation of large part of astro-specific genes. To ease the interpretation, we report the relative changes in cell type proportions as marker gene profiles (MGPs) selected by *markerGeneProfile* package (Figure 4A).

**Figure 4.**
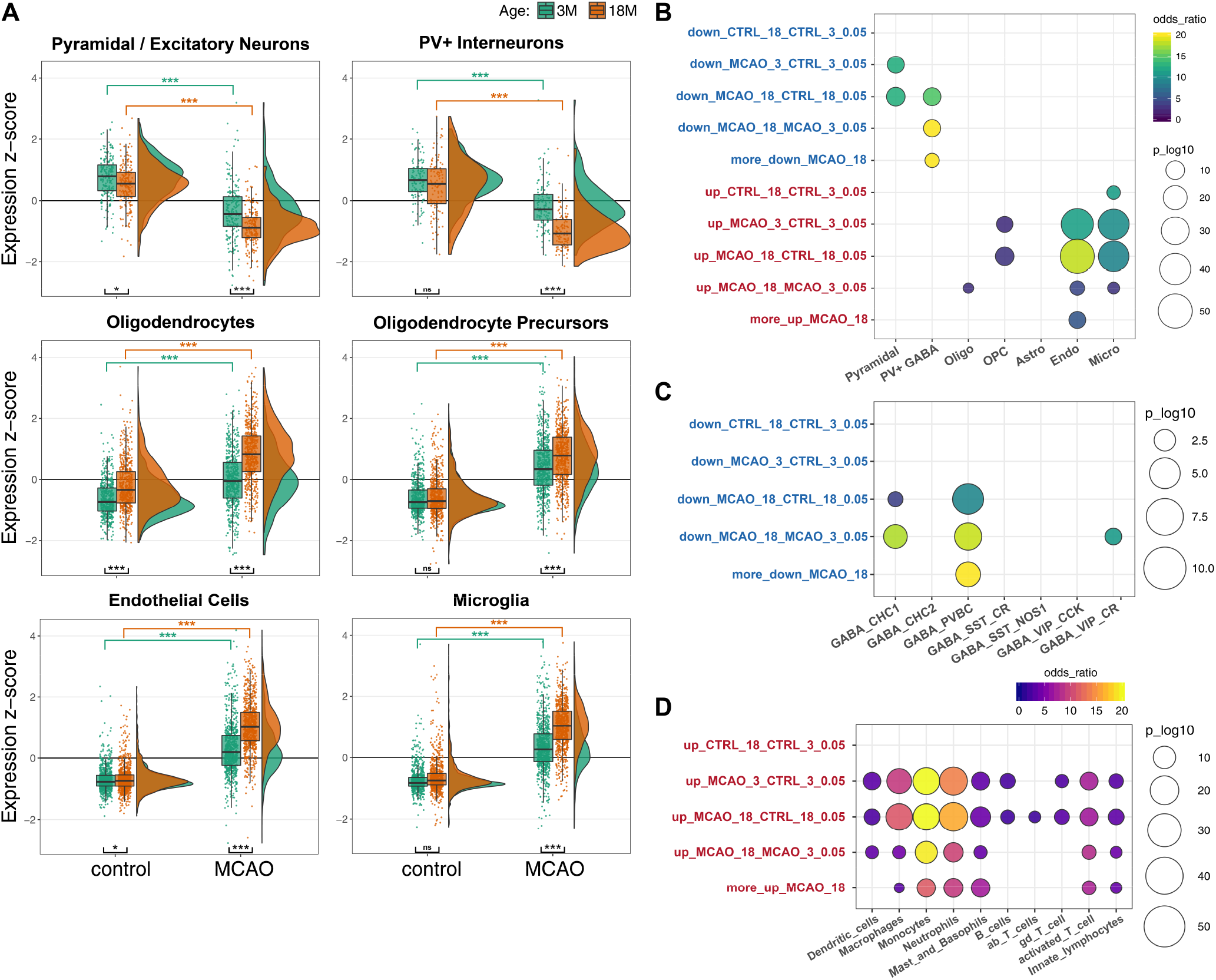
Cell type proportion estimation by transcriptome deconvolution. A) Raincloud plots showing z-scored expression of selected marker genes for major brain cell populations. Astrocytes not shown (see text and Figure S6). Asterisks show Holm-corrected post-hoc t-test p-values. Significance codes (***) <0.001; (**) <0.01; (*) <0.05; (ns) >0.05. B) Overlap between cell marker gene sets and the sets of differentially expressed genes. C) Overlap between sets of aging/stroke downregulated genes and the markers of seven phenotypically well-characterized GABAergic interneuron subpopulations. CHC = Chandelier cells; PVBC = PV+ fast-spiking basket cells; other subpopulations named after dominant markers (see 75). D) Overlap between sets of aging/stroke upregulated genes and the marker genes of the major leukocyte populations. See also Figures S7 and S8.

We found a significant increase in the MGPs of all non-neuronal cell types following stroke in both age groups (p_adj_ < 2.2e-16; Figure 4A). The largest increase was in microglial and endothelial marker genes (log_2_FC 1.26-1.77) and the lowest in oligodendrocytic markers (log_2_FC 0.47-0.60), which also significantly increased with normal aging (p = 1.20e-19; Figure 4A). Glial marker genes had generally higher expression in aged stroke group compared to young stroke group, although the magnitudes of their activation by stroke were relatively comparable between both ages. The most prominent difference was in endothelial cell markers (log_2_FC_young_ = 1.26, log_2_FC_aged_ = 1.74), which were also significantly overrepresented among “more up MCAO18” genes (Figure 4B). Furthermore, cell type proportion estimates revealed significant depletion of pyramidal/excitatory neurons during aging and following stroke in both age groups (p_adj_ < 1.00e-06, log_2_FC_young_ = −0.56, log_2_FC_aged_ = −0.63) without effect of interaction (p_interaction_ = 0.154; Figure 4A).

### Aged ischemic brain is characterized by selective vulnerability of PV+ interneurons and increased infiltration of peripheral leukocytes

Unlike pyramidal/excitatory neurons, parvalbumin-positive (PV+) GABAergic interneuron markers were down-regulated after stroke to a greater degree in aged animals (log2 FC young= −0.42, log_2_FC_aged_ = −0.89, p_interaction_ = 6.22e-08; Figure 4A), which was supported by the significant overlap with “more down MCAO18” gene set (Figure 4B), suggesting that this neuronal class is particularly vulnerable to ischemia in aged mice. PV+ interneurons play key roles in cortical plasticity and thus may have profound effect on post-stroke recovery^73,74^. Testing for enrichment of independent set of marker genes of 6 phenotypically well-defined interneuron populations^75^ also revealed significant overlap of “more down MCAO18” genes with markers of PV+ fast-spiking basket cells, but not other interneuron populations (Figure 4C). We then searched the literature and found that additionally, at least 30 out of 122 genes in the “more down MCAO18” gene set are directly linked to PV+ interneurons and are often localized in their projections^73,75,84–89,76–83^ (Figure S7A). We then confirmed the enrichment of these genes in PV+ interneurons using two recent single-cell RNA-Seq datasets^90,91^ (Figure S7B, C), providing strong indication that the decreasing transcriptional signal indeed reflects the selective impairment of PV+ interneurons in aged post-stroke mice. Afterwards we analyzed selected marker genes of CNS cell types at several time-points after stroke by RT-qPCR in a new set of mice (see below), which again revealed a significant impact of stroke on PV+ interneurons in aged (p = 6.95e-07), but not young mice (p = 0.156; Figure S8).

Since peripheral immune cells can infiltrate the brain following the disruption of blood-brain barrier after stroke, we assessed their contribution by analyzing the overlap of DE gene sets with cell-specific genes of several leukocyte populations (Figure 4D). Stroke-induced genes in both age groups were highly enriched with macrophage-, monocyte- and neutrophil-specific genes (p ≤ 4.19e-14) and less strongly with mast cell/basophil-, dendritic cell- and activated T-cell-specific genes (p ≤ 1.18e-5). Interestingly, granulocyte-enriched genes (neutrophils, mast cells/basophils) were significantly overrepresented among “more up MCAO18” DE genes (p ≤ 1.67e-6), as were monocyte-enriched genes (p = 2.14e-6) and, to a smaller degree, innate lymphocyte (NK cells; p = 0.025) and activated T-cell-enriched genes (p = 0.003), although T-cell pan markers were barely detectable in our sequencing data (Figure S9). These trends were recapitulated by expression of individual marker genes reported as highly specific and stable by at least two recent expression studies^59,60,92,93^ and by ImmGen database (www.immgen.org; Figure S9).

Overall, the analysis of relative cell type proportions highlighted both similarities and dissimilarities in cellular response to stroke between the two age groups. Similarities include proliferation of non-neuronal cells (particularly of microglia and endothelial cells), death of pyramidal neurons and temporal dynamics of cellular responses. Dissimilarities manifest in greater damage to PV+ interneurons, increased abundance of endothelial cells and increased infiltration of granulocytes/neutrophils in aged mice.

### Network analysis provides systems perspective on aging, stroke and their interaction

To capture the full extent of expression trends from systems perspective and reveal relationships between the genes, we complemented our results with weighted co-expression network analysis (WGCNA). We found 27 modules of highly co-expressed genes. Ten modules were strongly associated with aging and/or injury status, of which nine modules corresponded to stroke-induced or stroke-repressed genes (Figure S10A, B). The remaining module (darkturquoise) was upregulated with age, contained many oligodendrocyte-specific genes and was enriched in GO terms like “myelin sheath” and “metabolism of lipids” (Figures 5A-D, S10A), providing further support for positive correlation of oligodendrocyte-related expression with aging from previous analysis (Figure 4A). Among stroke-repressed modules, blue neuron-associated module was similarly downregulated in both young and aged mice, while green module showed greater downregulation in aged mice (Figure 5C). The green module was significantly enriched in markers of PV+ interneurons (p = 9.29e-06) and synaptic-transmission- and axon-related genes, showing that selective axonal damage of PV+ interneurons is embodied at the system-wide transcriptional level (Figure 5B, D). The turquoise module contained a large number of genes upregulated after stroke in both age groups, majority of which were immune/inflammation- and cell cycle-related and was significantly enriched with microglial and endothelial markers (Figure 5A-D). This module likely reflects the increased proportion and activation of these cell types after stroke.

**Figure 5.**
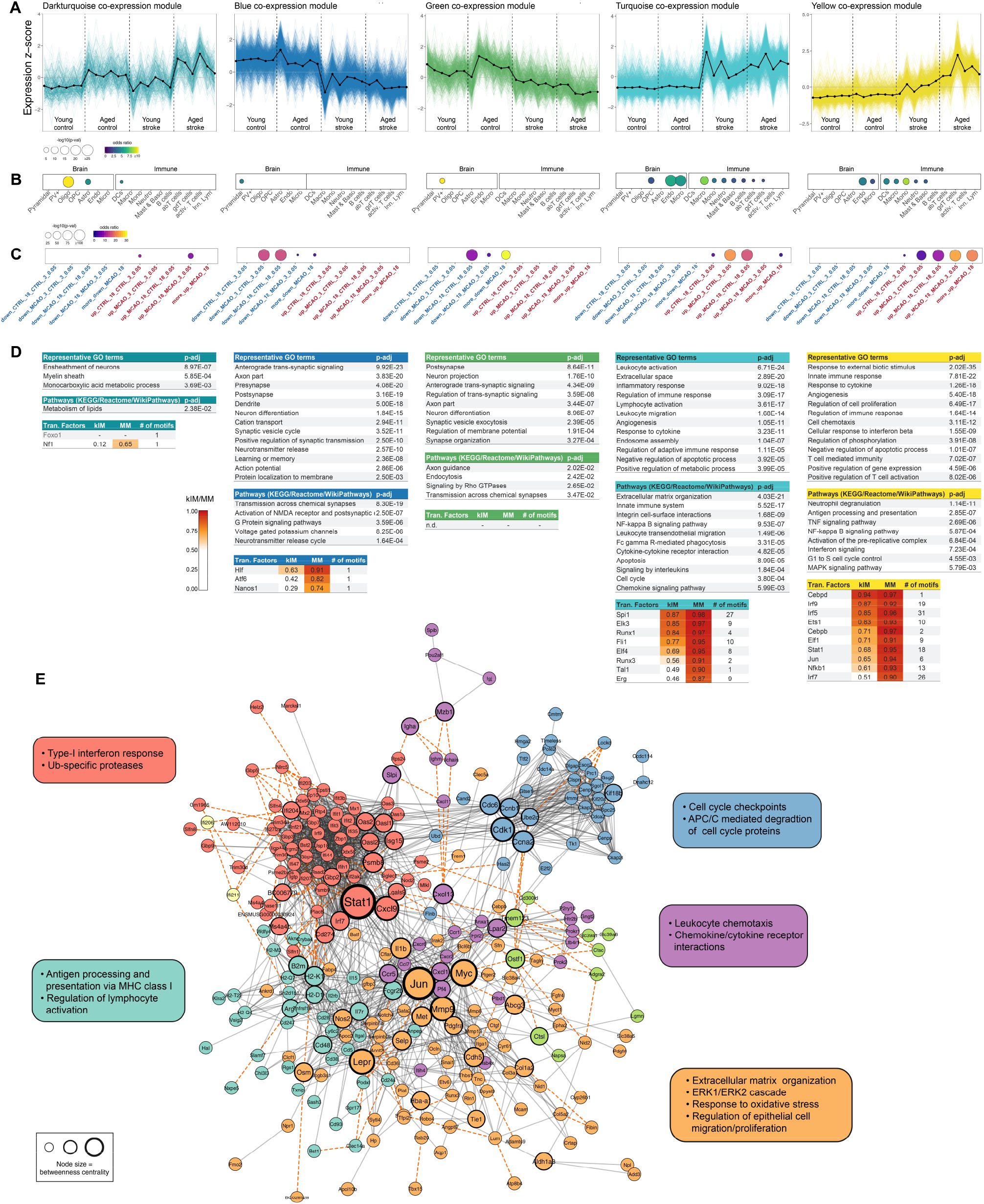
Gene co-expression and interaction network analysis. A) Expression of selected WGCNA modules associated with aging and/or stroke. B) Module enrichment with cell-specific genes of major CNS and immune cell populations. C) Module enrichment with differentially expressed genes. D) Functional characterization of modules via enrichment of gene ontology sets, cellular pathways and targets of transcription factors. kIM = intramodular connectivity. MM = module membership. E) Protein interaction network constructed from “more up MCAO18” DE gene set. Network sub-clusters with functional annotations are shown in different colors. Key signaling hubs with high betweenness centrality highlighted with increasing node size. See also Figure S10.

In addition, turquoise genes were significantly enriched with targets of transcription factors such as *Spi1, Runx1, Runx3, Tal1, Fli1, Elf4* - known master regulators of hematopoietic development^94,95^ and microglial homeostasis and activation^96,97^ (Figure 5D). The yellow co-expression module was more up-regulated by stroke in aged mice compared to young mice, significantly enriched in leukocyte and endothelial-specific genes, with many GO terms related to inflammation, cytokine production, antigen processing and presentation, leukocyte chemotaxis, lymphocyte activation and interferon signaling as well as regulation of gene expression, suggesting that the yellow module is largely subject to intrinsic transcriptional activation (Figure 5A-D). Supporting this, in the yellow module we detected the largest number of enriched transcription factors (relative to other modules) including *Stat1, Cebpd, Cebpb, Ets1, Elf1, Jun, Nfkb1* and many interferon regulatory factors (*Irf9, Irf5, Irf7*) (Figure 5D). In accordance, there was a striking overrepresentation of interferon-stimulated genes (p = 1.25e-50). Together, WGCNA largely recapitulated results of cell type proportion estimates in an unsupervised manner, supported our previous observation that no neuroprotective transcriptional program is activated specifically in young animals (as no such module was detected by WGCNA), and highlighted amplified activation of yellow inflammatory/interferon module by stroke in aged mice.

We noted that yellow co-expression module generally corresponds to the “more up MCAO18” DE gene set. Vast majority of the “more up MCAO18” DE genes either directly belonged to, or was highly correlated with the yellow module (Figure S10C, D). To investigate how these genes interact, we constructed a combined protein interaction network (Figure 5E). 227 out of 409 genes in the “more up MCAO18” DE set were highly connected within the network. Network clustering revealed six clusters enriched in related, but distinct functional terms including cluster of interferon-stimulated genes, cluster of genes involved in antigen presentation, cluster of cell-cycle regulatory genes, chemokine/cytokine and chemokine receptors cluster and one less rigid cluster of genes involved in extracellular matrix organization, ERK/MAPK signaling and response to oxidative stress (Figure 5E). Searching the network for signaling hubs with high betweenness centrality highlighted transcription factors *Stat1, Jun* and *Myc* as well as matrix metalloproteinase 9 (*Mmp9*), chemokine ligands *Cxcl9* and *Cxcl13*, leptin receptor (*Lepr*), and cyclin-dependent kinase 1 (*Cdk1*) as key hubs acting as crosstalks between functional clusters of the network (Figure 5E). These results suggest that “more up MCAO18” genes are part of the transcriptional program composed of several distinct, but molecularly connected gene modules.

### Age-dependent activation of type-I IFN regulatory modules after stroke

A striking hallmark of the differential response of aged animals to ischemic stroke was activation of type-I interferon signaling pathway (Figures 2B, 3A, 5A-E, S2, S3B). Type I interferons (IFN-Is) are key antiviral cytokines that elicit prototypical interferon-stimulated genes (ISGs) encoding antiviral and inflammatory mediators^98^ and also activate other signaling pathways including MAPK cascades, other cytokines, chemokines and cell-cell interaction modifiers (MHC-I, *Lgals9)*^99^. It has been reported that blocking the IFN-I signaling improves stroke outcome in young mice^100^. IFN-I signaling may therefore act as the central player in the increased neuroinflammatory signature that we detected in the aged animals post-stroke, aggravating the neuronal injury.

To explore this signaling component in greater detail, we mapped our expression data onto the recently published cross-species IFN-I regulatory network^99^ (Figure 6B-D). IFN-I network is divided into five regulatory clusters (C1-C5) with functional differences and variable disease associations^99^. Mapping our data against the network revealed that cluster C3 (composed mainly of antiviral effector genes) and C5 (enriched in inflammation mediators and regulators) were upregulated after stroke in both aged and young animals while the remaining three clusters were non-responsive (Figure 6B). Of the two responsive clusters only cluster C3 was differentially induced between young and aged animals (p = 3.63e-36). C3 represents a putative cluster containing predominantly genes under control of ISGF3 transcription factor complex composed of STAT1, STAT2 and IRF9^99^. In accordance, several regulators of the C3 cluster including components of ISGF3 complex were part of “more up MCAO18” DE set (Figure 6C, D); altogether suggesting that age-dependent amplification of ISG expression occurs in a STAT-dependent manner.

**Figure 6.**
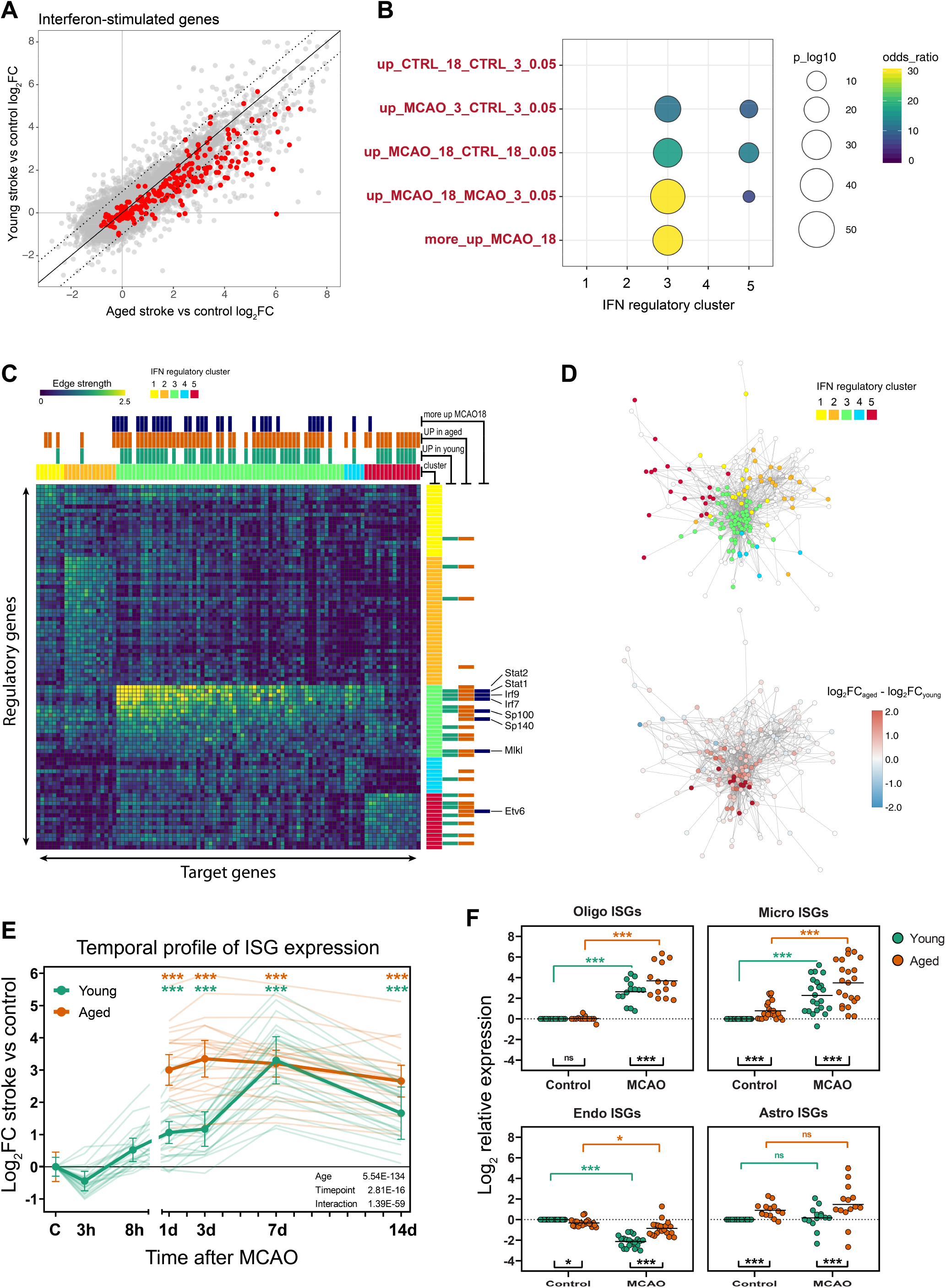
Transcriptional response of IFN-I signaling after stroke in young and aged mice. A) Scatter plot of stroke-induced log_2_ fold-change in young and aged mice. Highlighted is a set of 207 interferon-stimulated genes. B) Enrichment of IFN-I regulatory network clusters from 99 in aging/stroke upregulated gene sets. C) Heatmap of IFN-I network regulatory links between regulators (kinases, phosphatases, transcription factors) and target genes reconstructed from 99. Binarized mapping of genes to stroke-upregulated gene sets in young/aged mice is shown on top and right. D) Network visualization of IFN-I regulatory network; 1000 strongest edges are shown. Network is colored by regulatory cluster (top) or by difference in stroke-induced log_2_FC between aged and young mice (bottom). E) Time-course analysis of ISG expression after stroke in young and aged mice. Bold line shows average expression of 23 ISGs measured by RT-qPCR. Error bars show standard deviation of biological replicates (n = 2-5). Thin lines show average expression of each gene. Asterisk show significant difference relative to control for each age separately (mixed model with post-hoc t-tests). Mind the break in x-axis. Due to limited number of 18-month old animals, 3h and 8h time points were omitted for the aged group. See also Figure S11 for expression of additional genes and Table S2 for per-gene statistics. F) ISG expression in purified cell populations 3 days after MCAO from young and aged mice. Asterisks show Holm-corrected post-hoc t-test p-values. See also Figure S12. G) Significance codes: (***) <0.001; (**) <0.01; (*) <0.05; (ns) >0.05.

### Post-stroke temporal dynamics of IFN-I signaling in young and aged mice

IFN-I signaling is typically characterized by rapid ISG induction, which is afterwards quickly attenuated by negative feedback mechanisms^99^. As the outcomes of the IFN-I signaling within the CNS may depend on a delicate balance^98,101^, we asked if the increased IFN-I signature in the aged post-stroke mice is due to constitutively greater expression or rather altered expression dynamics relative to young mice. To answer this question, we analyzed the expression of IFN-Is, IFN receptors, ISGs and other selected genes by microfluidic RT-qPCR at several time points following stroke (Figures 6E, S11).

As expected, all 23 ISGs changed in a largely coordinated fashion throughout the time course, albeit with varying fold-changes (Figure 6E). Expression of ISGs in young animals initially slightly decreased at 3h post MCAO and then started to increase, with peak at 7 days after MCAO (average log_2_FC= 3.30; n = 23 ISGs). In aged mice, the increase of ISG levels was on average four times greater at day 1 and day 3 compared to the increase in young animals. Unlike in young mice, the expression of ISGs did not further fluctuate, but remained at elevated levels until the latest investigated time point (14 days, average log_2_FC = 2.66). Together, these results show that activation of IFN-I signaling occurs early, but not sooner than 3h after stroke, and both timing and overall magnitude of ISG expression are important factors in the different response of aged brain to ischemic injury.

### Cell-specific analysis of IFN-I signaling in young and aged mice after stroke

*Stat1* activation in neurons^102^ and more recently increased IFN-I signaling in microglia^103^ following tMCAO have been demonstrated in young animals. However, little is known about the IFN-I signaling in other cell types following ischemic stroke, and no cell type specific data are available in the aged brain. We have therefore focused on the key players in the brain inflammatory responses – glia and endothelial cells^104^ – and aimed to identify how they contribute to IFN-I signaling following stroke. We have FACS-sorted populations of astrocytes (GFAP+), microglia (CD11b+), oligodendrocytes (O4+) and endothelial cells (CD31+) from young and aged mice 3 days after MCAO as well as from the age-matched controls and measured expression of ISGs and other selected genes by microfluidic RT-qPCR (Figure 6F, S12).

Microglia and oligodendrocytes heavily upregulated ISG expression following stroke in both young and aged animals (Figures 6F, S12B). Endothelial cells displayed opposite behavior and significantly downregulated vast majority of measured ISGs in young, and to a much lesser extent in the aged mice. In astrocytes, 9 out of 23 ISGs were undetected. The expression of the remaining 14 ISGs did not change in a synchronized fashion, altogether showing a limited response of astrocytes to IFN after stroke in both age groups. In addition, astrocytes and microglia, but not endothelial cells and oligodendrocytes, showed significant ISG upregulation with normal aging (Figure 6F).

Next, we assessed the relative differences in ISG expression between cell types for each experimental group separately (Figure S12C). This comparison revealed that in young and aged controls, endothelial cells are the main ISG expressors, both in terms of number of expressed ISGs and their expression levels, suggesting homeostatic role of IFN-I signaling in these cells. In contrast, microglia, oligodendrocytes and astrocytes expressed lower ISG levels under homeostasis and maintained predominantly expression of ISGs with regulatory roles, such as transcription factors (*Stat1, Stat2, Irf9*) and receptors (*Ifih1, Ddx58*). After stroke, the relative cell type contributions changed and microglia expressed similar levels of ISGs to endothelial cells (Figure S12C). Overall, our cell-specific analysis revealed that not only microglia, but also oligodendrocytes heavily induce IFNI signaling following stroke. All cell types converge on higher ISG expression in aged post-stroke brain, although their individual contribution and response to stroke differ. Despite this trend, it is likely that effects in bulk tissue reflect also ISG induction in other cell types, such as non-microglial immune cells entering the brain through compromised BBB.

## Discussion

In this study, we systematically analyzed the impact of aging, stroke and their interaction on genome-wide expression profiles. Several findings emerged from the analysis, including that i) brain aging is accompanied by increased inflammation driven by alterations of glial cells, ii) transcriptional response to stroke in young and aged brain is highly similar and differs primarily in magnitude, iii) aged PV+ GABAergic interneurons are particularly vulnerable to stroke with potential implications for functional recovery, iv) differential stroke outcome is associated with over-activation of pro-inflammatory pathways and other potentially detrimental factors in aged mice, rather than activation of neuroprotective program in young mice, v) increased activation of IFN-I signaling represents a key difference in response to stroke between young and aged mice vi) subnetworks of IFN-I signaling are differentially sensitive to combination of stroke and aging, vii) temporal profile of IFN-I activity after stroke differs between young and age mice and viii) microglia and oligodendrocytes, but not astrocytes and endothelial cells massively upregulate IFN-I signature following stroke.

First, we analyzed factors that may sensitize aged brain to greater stroke damage during normal aging process. A number of studies has documented that inflammation is increased in the aged brain^105–107^, and it has been reported that the changes occur predominantly in glial cells^108,109^. Our results are well in line with these reports as we detected upregulation of a number of pro-inflammatory genes (Figures 1A, S1A) that map to signatures of aged and reactive glial cells, including the signature of a recently identified subpopulation of *Ccl4*+ pro-inflammatory microglia^60,61^ (Figures 1B, S2). Activated microglia can promote astrocyte reactivity through secretion of modulatory factors^110^, and thus further amplify the neurotoxic environment. Together, these changes are likely to sensitize aged brain towards more severe stroke and secondary injury. In addition, we detected increase in oligodendrocyte-specific expression with normal aging (Figures 4A, 5), which has been also seen in human brain^111,112^ and can be possibly attributed to increased number of oligodendrocytes^113–115^.

Next, we investigated how adult and aged brain responds to ischemic challenge. The assessment of cellular differences by computational deconvolution revealed that PV+ GABAergic interneurons are particularly vulnerable to ischemic stroke in aged mice (Figures 4A-C, S8). This result was recapitulated by RT-qPCR (Figure S8) and unsupervised WGCNA analysis where we detected PV+ interneuron-associated module (“green") with greater repression in aged post-stroke mice (Figure 5). Dysfunction or loss of PV+ interneurons is implicated in the pathology of numerous neuropsychiatric disorders, including schizophrenia^116,117^, Alzheimer’s disease^118^ and depression^119,120^. Given their central role in the modulation of neuronal plasticity and cortical information processing^73,74,121,122^ – processes that underlie recovery after stroke^123^ – the PV+ interneurons may represent novel therapeutic target to promote functional recovery in elderly stroke patients.

Surprisingly, we did not detect any exclusive or greater activation of neuroprotective genes in young mice compared to aged mice (Figure 2). Instead, stroke in young mice more highly induced similar pro-inflammatory signatures of activated glial cells that are already upregulated with normal aging (Figures 2, S2). On the other hand, in aged post-stroke mice we found markedly stronger upregulation of >400 genes, many involved in the processes of the inflammatory cascade (Figure 3A). This was accompanied by a greater influx of peripheral leukocytes, particularly neutrophils (Figure 4D), which are the strongest producers of reactive oxygen species (ROS) and matrix metallopeptidases (MMPs) and promote neuronal injury and BBB disruption^124–126^. In agreement, an increased number of neutrophils with altered phenotypic responses was previously seen in aged male mice and human stroke patients compared to their younger counterparts^127^ and an increased neutrophil to lymphocyte ratio has been associated with stroke severity and outcome^128^. Although the absence of neuroprotective signal in bulk tissue does not rule out the possibility of its presence in individual cell types, these results suggest that an increased neuroinflammation and infiltration of circulating immune cells are one of the primary drivers for the exacerbated pathology in aged mice.

An outstanding feature of differential response of aged animals to ischemic stroke was upregulation of IFN-I pathway (Figure 2B, 3A, 5A-E, S2, S3B), which persisted for at least 14 days (Figure 6E). IFN-Is are antiviral cytokines with pleiotropic roles^129^ implicated in a number of CNS pathologies including multiple sclerosis^130,131^, Aicardi-Goutières syndrome, amyotrophic lateral sclerosis^132^, Alzheimer’s disease^59,133,134^, spinal cord injury^135^, traumatic brain injury^136,137^ and ischemic stroke100,103,138.

Therefore, we have explored INF-I signaling in more detail and found that two IFN-I regulatory modules are activated by stroke irrespective of age, but only the canonical STAT-dependent module is differentially activated in aged animals (Figure 6B-D), likely contributing to an increased neurotoxicity^100^. Indeed, *Stat1* and *Irf9* have deleterious roles in stroke and can act directly on neurons^102,138^. ISG activation typically requires both IFNAR1 and IFNAR2 receptor subunits^139^. Recently the IFNAR2- and STAT-independent pathway, triggered by IFN•, was shown to be involved in systemic LPS-induced toxicity^140^. While the genetic or pharmacological blocking of IFNAR1 leads to neuroprotection after transient MCAO^100^, the same study reported no effect in IFNAR2−/− mice. This suggests the involvement of the compensatory IFNAR2- and STAT-independent pathway after stroke as well, although it has not been explored in aged animals. Our finding of a predominant increase in the IFNAR2- and STAT-dependent module in aged animals indicates that the detrimental effects of IFN-I signaling after stroke may be exerted by ISGs that are common to the IFNAR2-/STAT-dependent and independent pathways.

As the cell-specific context to the IFN-I signaling after stroke occurance has not been well described in the literature, we profiled the responses of the glia and endothelial cells – known key players in brain neuroinflammatory responses^104^ (Figures 6, S12). Our results support the perspective that not only microglia, but also oligodendrocytes are active players in the acute inflammatory response after stroke. On the other hand, although endothelial cells expressed the highest levels of ISGs under control conditions, they downregulated IFN-I pathway after stroke. This disparity may be associated with different roles of IFN-I signaling in the endothelial cells, evident by the vastly different baseline ISG levels, in line with the role IFNb plays in the maintenance of the BBB integrity^98^.

Despite the overall trend of increased ISG expression in the aged post-stroke brain in the analyzed cell populations, it is unlikely to explain alone the overall ISG increase we reproducibly detected in the bulk tissue. These results indicate the involvement of other contributors, such as the early-infiltrating peripheral leukocytes. Previously, the hematopoietic component has been identified as a major source of IFN-I signaling following traumatic brain injury^136^ and aged mice with bone-marrow transplants from young mice have improved stroke outcome^127^. In concordance, we detected greater upregulation of granulocyte signature genes in aged animals (Figure 4D). In addition, it has been suggested that IFN levels correlate with severity of injury and differently influence functional outcome, as IFN signaling is beneficial in context of mild ischemic preconditioning^141,142^, but it is detrimental following more severe stroke^100^. Our findings appear to be consistent with this notion, as aged animals generally suffer more severe strokes. One potential mechanism of this phenomenon could be the greater disruption of BBB, which in turn leads to the greater influx of peripheral leukocytes. Nonetheless, it is clear that the differential activation of IFN-I signaling pathway in aged animals is likely to contribute significantly to the exacerbated stroke outcome in aged mice and represents a potential target for therapeutic intervention that has been so far overlooked. Our results provide one of the first steps in this direction and open the door to future studies needed to address the mechanisms underlying IFN-I neurotoxicity following stroke in the aged brain.

As with the majority of studies, our results also need to be viewed in light of potential limitations. Since our RNA-Seq data are based on bulk tissue, the expression signal is partly confounded by the cell type composition and the power to detect genes altered in a cell-specific manner is lowered. To tackle this effect, we have employed cell type deconvolution techniques and assayed the results with a range of cell-specific signatures. Another limitation is that our RNA-Seq experiment assayed a single time-point (3 days) after experimental stroke relative to the control. Although the selected time-point can be considered well representative of the subacute phase, informative on early damage as well as initiation of repair processes^143,144^, there is a space for future studies analyzing later subacute and chronic phases. Being aware of the aforementioned limitations, we provide direct cell-specific as well as time course data targeted at the most significant findings.

In conclusion, detailed insights into transcriptional response to stroke described in this study may contribute to our understanding of the interplay between stroke pathology and aging, and open new avenues for the future search for effective therapeutic approaches.

## Methods

### Animals

Experiments were performed on 3 and 18 month-old C57Black/6 or FVB female mice. FVB mice are GFAP/EGFP transgenic (line designation TgN(GFAP-EGFP), FVB background), in which the expression of enhanced green fluorescent protein (EGFP) is controlled by the human glial fibrillary acidic protein (GFAP) promoter (marker of astrocytes)^145^. The mice were kept on a 12-hr light/dark cycle with access to food and water *ad libitum*. All procedures involving the use of laboratory animals were performed in accordance with the European Communities Council Directive 24 November 1986 (86/609/EEC) and animal care guidelines approved by the Institute of Experimental Medicine, Academy of Sciences of the Czech Republic (Animal Care Committee on April 7, 2011; approval number 018/2011). All efforts were made to minimize both the suffering and the number of animals used.

### Induction of middle cerebral artery occlusion (MCAO)

Prior to the induction of MCAO, mice were anaesthetized with 3% isoflurane (Abbot) and maintained in 2% isoflurane using a vaporizer (Tec-3, Cyprane Ltd.). A skin incision between the orbit and the external auditory meatus was made, and a 1-2 mm hole was drilled through the frontal bone 1 mm rostral to the fusion of the zygoma and the squamosal bone and about 3.5 mm ventrally to the dorsal surface of the brain. The middle cerebral artery (MCA) was exposed after the dura was opened and removed. The MCA was occluded by short coagulation with bipolar tweezers (SMT) at a proximal location, followed by transection of the vessel to ensure permanent occlusion. During the surgery, body temperature was maintained at 37±1°C using a heating pad. This MCAO model yields small infarct lesions in the parietal cortical region. Intact cortical tissue from 3 and 18 month-old mice was used as control.

### Dissection of brain tissue from the mouse cortex

Mice were deeply anesthetized with pentobarbital (PTB) (100 mg/kg, i.p.), and perfused transcardially with cold (4–8°C) isolation buffer containing (in mM): NaCl 136.0, KCl 5.4, Hepes 10.0, glucose 5.5, osmolality 290 ± 3 mOsmol/kg. To isolate the cerebral cortex, the brain (+2 mm to −2 mm from bregma) was sliced into 600 μm coronal sections using a vibrating microtome HM650V (MICROM International GmbH), and the uninjured or post-ischemic parietal cortex was carefully dissected out from the ventral white matter tracks.

### Preparation of single cell suspensions

The collected tissues were incubated with continuous shaking at 37°C for 45 min in 2 ml of papain solution (20 U/ml) and 0.2 ml DNase (both Worthington) prepared in isolation buffer. After papain treatment, the tissue was mechanically dissociated by gentle trituration using a 1 ml pipette. Dissociated cells were layered on the top of 5 ml of Ovomucoid inhibitor solution (Worthington) and harvested by centrifugation (140 x g for 6 min). This method routinely yielded ~2 x 10^6^ cells per mouse brain. Cell aggregates were removed by filtering with 30 μm cell strainers (Becton Dickinson). The cell suspension was then labeled for oligodendrocyte marker 1:50 anti-O4-PE (Miltenyi Biotec), endhothelial cell marker 1:50 anti-CD31-PE (Miltenyi Biotec) and microglial marker 1:50 anti-CD11b-APC (Miltenyi Biotec), according to the standard manufacturer’s protocol. To collect astrocytes, GFAP/EGFP mice were used. The cells were kept on ice until sorting.

### Cell sorting and collection

Single cell suspensions were sorted using flow cytometry (FACS; BD Influx). Hoechst 33258 (Life Technologies, Carlsbad, CA, USA) was added to the suspension to check viability. 100 cells per well were collected into 96-well plates (Life Technologies) containing 5 μl nuclease-free water with bovine serum albumin (1 mg/μl, Fermentas) and RNaseOut 20 U (Life Technologies). The plates were placed on a precooled rack and stored at −80 °C until analysis. After discarding samples of insufficient quality, we used 2-5 mice per cell type per experimental group for the RT-qPCR analysis (see below). We note that all samples were strongly enriched in respective cellular markers and depleted in markers of other cell types, confirming high purity of sorted suspensions (Figure S12A).

### RNA isolation, library preparation and sequencing

Brain tissue samples were homogenized using the Tissue-Lyser (Qiagen). Total RNA was extracted with TRI Reagent (Sigma-Aldrich) according to the manufacturer’s protocol and treated with TURBO DNA-free kit (Thermo Fisher). RNA quantity and purity was assessed using the NanoDrop 2000 spectrophotometer (Thermo Fisher) and RNA integrity was assessed using the Fragment Analyzer (Agilent). All samples had RQN > 8. Libraries were prepared from 400 ng total RNA with QuantSeq 3’ Library Prep Kit FWD (Lexogen) according to manufacturer’s protocol. 1 μl of ERCC spike-in (c = 0.01x; Thermo Fisher) per library was included. This library preparation method generates stranded libraries predominantly covering the 3’ end of the transcript, thus producing genecentric expression values. Libraries were quantified on the Qubit 2 fluorometer (Thermo Fisher) and Fragment Analyzer (Agilent) and sequenced on the NextSeq 500 high-output (Illumina) with 85 bp single-end reads. 11.5 – 38 million reads were obtained per library with a median of 16 million reads.

### RNA-Seq data processing, mapping and counting

Adaptor sequences and low quality reads were removed using TrimmomaticSE v0.36^146^. Reads mapping to mtDNA and rRNA were filtered out using SortMeRNA v2.1 with default parameters^147^. The remaining reads were aligned to GRCm38 and ERCC reference using STAR v2.5.2b with default parameters^148^. Mapped reads were counted over Gencode vM8 gene annotation using htseq-count with union mode for handling of overlapping reads^149^.

### Differential expression analysis

Several comparisons of differential expression were generated using DESeq2 v1.16.1^150^. For pairwise comparisons, we compared aged controls to young controls, young stroke group to young controls, aged stroke group to aged controls and aged stroke group to young stroke group (p_adj_ < 0.05, log_2_FC >1 for upregulation and < −0.65 for downregulation). We also generated two-factor comparisons using injury (control/MCAO) and age (3m/18m) and their interaction as predictor variables. DESeq2 results are available in Supplementary file 1. Searchable visualization table is available in Supplementary file 2. During the initial analysis of the dataset, we noted that there was a relatively large number of genes induced or repressed exclusively, or with a greater fold-change in aged animals, although only a subset of them reached statistically significant interaction term as outputted by DESeq2 analysis. In order to identify all of the genes that are likely subject to age-stroke interaction, we have prepared four additional sets of genes with age-dependent differential response to stroke containing genes that are significantly influenced by stroke in one age group and at the same time their fold-change (vs control) is at least doubled compared to second age group. That is, the set “more up MCAO18” is comprised of genes significantly upregulated in aged animals after stroke (compared to aged controls; p_adj_ < 0.01, log_2_FC_aged_ > 1), and at the same time having significant interaction term (p_adj_ < 0.1) and/or having at least double the fold change of young strokes (compared to young controls; log_2_FC_aged_ − log_2_FC_young_ > 1). The same rationale was applied for more highly upregulated genes in the young stroke group (“more up MCAO3”) and for downregulated genes in both age groups (“more down MCAO18”, “more down MCAO3”), with an exception that the log_2_FC threshold was < −0.65 for downregulation. DE sets legend is also available in Table S1. Gene sets composition can be found in Supplementary file 3. Only genes with average expression ≥ 5 normalized counts in at least one experimental condition were considered for further analysis.

### Gene set enrichment analysis (GSEA)

GSEA^50^ was performed for pairwise differential expression comparisons. First, a gene score was calculated for every gene using DESeq2 output as −log_10_(p_adj_) and assigned a positive or negative sign based on direction of regulation. Genes were ranked by their gene-scores and GSEA was run in a weighted pre-ranked mode with 1000 permutations. Gene sets were downloaded from http://download.baderlab.org/EM_Genesets/, and GSEA was run separately for two gene ontology (GO) categories (biological process – GOBP, cellular component – GOCC). Only gene sets containing between 15 and 1000 (for GOBP) or 5 to 1000 (for GOCC) genes were considered. Annotations with IEA (inferred from electronic annotation) evidence codes were excluded. For pathway enrichment, a gene set file integrating several pathway databases was used. Significantly overrepresented gene sets were visualized as a network using Enrichment Map ^62^. In the network, each node represents gene set and highly overlapping gene sets are connected with edges, resulting in a tight clustering of highly redundant gene sets. For functional annotation of discrete sets of genes we used Cytoscape plugin ClueGO^151^ with the following parameters: no IEA codes, right-sided hypergeometric test with Benjamini-Hochberg correction for statistical testing and all genes after filtering (16048 genes) as the background set.

### Cell-specific gene sets and cell type proportion estimation

Marker genes for major cell types specifically in the mouse cortex region were taken as an initial reference^71^ (Supplementary File 3). Unlike other marker databases that rely on a single data source, this marker set represents a consensus from several published studies and accounts for brain regional heterogeneity. The microglial marker genes in the reference marker set were already devoid of genes differentially expressed in activated microglia^53^. In order to acquire marker genes with stable expression regardless of activation states, we have further removed the genes previously found to be differentially expressed in microglia after tMCAO^152^; in aged cortical microglia^54^, and aged whole-brain microglia^53^ compared to young microglia; and genes enriched in bone marrow-derived macrophages compared to microglia^153^. We have also removed genes that were differentially expressed in astrocytes after tMCAO^64^ and aged cortical astrocytes^51,52^. Because peripheral immune cells may infiltrate the brain following stroke, we have also excluded genes enriched in the major leukocyte populations obtained from ImmGen database (www.immgen.org).

DESeq2-normalized gene expression data and the cell-specific gene lists were used as an input into the *marker-GeneProfile* R package v1.0.3^71^ for the estimation of marker gene profiles (MGPs), which serve as a proxy for relative cell type proportion changes. We used a more stringent expression cutoff (average • 5 normalized counts across all samples) to reduce the transcriptional noise. Since it still may be possible that some marker genes are transcriptionally regulated under our experimental conditions, genes with reduced correlation (potentially regulated) to the majority of marker genes (assumed to reflect primarily cell type proportion change) were excluded from final estimates as described in^71^. Resulting MGP estimates were flagged if a high proportion of marker genes was removed in the previous step (> 40%) and/or proportion of variance explained by the first principal component was low (< 50%). Differences in expression of final marker gene sets were analyzed by linear mixed model in R project v3.6.0 using *lmerTest* package v3.1^154^ on log_2_ transformed, DESeq2-normalized expression values. We used two-factor design (age, injury) with interaction as fixed effects and gene intercept as random effect. Significance was tested by Satterthwaite’s method. Post-hoc t-tests were performed using *emmeans* package v1.3.5 and p-values were adjusted by Holm method.

To validate the first estimations, we employed CIBERSORT, a transcriptome deconvolution algorithm that uses gene expression matrix of individual cell types as a reference, and deconvolutes the cellular composition of mixed sample by linear support vector regression^72^. We used published single-cell RNA-Seq dataset of adult mouse cortex as a reference gene expression signature^155^. From the normalized gene-expression matrix, we excluded all intermediate cells as definded by authors and used the remaining 1424 core cells assigned to the major CNS cell types. CIBERSORT was run in the relative mode with 1000 permutations, quantile normalization was disabled as recommended for RNA-Seq data and q-value cutoff was lowered to 0.15. The results were correlated to the corresponding cellular MGPs (Figure S6).

### Weighted gene co-expression network analysis (WGCNA)

Standard WGCNA procedure was followed to create gene co-expression networks using blockwiseModules function from the *WGCNA* R package v1.68^156^. Filtered DESeq2-normalized expression data were used to calculate Pearson correlation between all gene pairs. Soft-thresholding was then applied by raising correlation values to a power of 22 to amplify disparity between strong and weak correlations. The soft-thresholding power was chosen to achieve approximately scale-free network topology, as recommended for biological networks^157,158^.

The resulting signed adjacency matrix was used to calculate topological overlap matrix (TO), which was then hierarchically clustered with (1-TO) as a distance measure. Genes were then assigned into co-expression modules by dynamic tree cutting algorithm requiring minimal module size of 20 genes. Modules with a distance between the module eigengenes (MEs) of less than 0.2 were merged. ME is the first principal component of the gene expression values within a module and is used to summarize the module’s expression. Pearson correlation between each gene and ME was then calculated. This value is called module membership and represents how close a particular gene is to a module. Finally, each gene was assigned to a module for which it had the highest module membership.

### Motif and transcription factor enrichment analysis

Cytoscape plugin iRegulon^159^ with default parameters was used to search for over-represented motifs and their associated transcription factors 500 bp upstream of the transcription start site. All genes from a particular WGCNA module were used as an input. A transcription factor was considered a hit for a given module only if its gene belonged to the same module.

### Protein-protein interaction network

Known interactions (minimal interaction score 0.4) between genes in the “more up MCAO18” DE gene set were downloaded from STRING database v10.5^160^. Remaining unconnected genes from “more up MCAO18” gene set were then added to the network based on their correlation with any of the genes already present in the network requiring Pearson r ≥ 0.96 (edges visualized with orange dotted line in Figure 5E). The resulting interaction network was then visualized and analyzed in Cytoscape v3.5.1. Spectral partition-based network clustering algorithm^161^ via Cytoscape ReactomeFI plugin v6.1.0^162^ was used for network clustering.

### Custom gene set enrichment

Gene sets of interest were collected directly from relevant publications. For references and legends, see Table S1. R package *GeneOverlap* v1.20 was used to calculate the odds ratio (OR) and the significance of the overlap of the gene sets of interest with the Fisher’s exact test.

### High-throughput RT-qPCR

Samples were reverse transcribed in a reaction volume of 10 μl containing: 5 μl template (either 125 ng total tissue RNA or 100 sorted cells after direct lysis), 0.5 μl spike-in RNA (Tataa Biocenter; c = 0.1x for tissues or 0.01x for sorted cells), 0.5 μl equimolar mixture of random hexamers with oligo(dT) (c = 50 μM), 0.5 μl dNTPs (c = 10 mM), 2 μl 5× RT buffer, 0.5 μl RNaseOUT, 0.5 μl Maxima H-Reverse Transcriptase (all Thermo Fisher) and 0.5 μl nuclease-free water. After the pre-incubation step at 65°C (t = 5 min), followed by the immediate cooling on ice, the main incubation was performed at 25°C (t = 10 min), 50°C (t = 30 min), 85°C (t = 5 min), after which the samples were immediately cooled on ice. cDNA from tissue samples was diluted 4x in nuclease-free water; sorted cell-cDNA was left undiluted. All cDNA samples were pre-amplified immediately after reverse transcription in 40 μl total reaction volume containing 4 μl cDNA, 20 μl IQ Supermix buffer (Bio-Rad), 4 μl primer mix of 96 assays (c = 250 nM each), and 12 μl of nuclease-free water. Reactions were incubated at 95°C (t = 3 min) following by 18 cycles of 95°C (t = 20 s), 57°C (t = 4 min) and 72°C (t = 20 s). After thermal cycling, reactions were immediately cooled on ice and diluted in nuclease-free water (sorted cells 4x, tissue 50x). High-throughput qPCR was then performed on a 96.96 microfluidic platform BioMark (Fluidigm) as previously described^163^. Cycling program consisted of activation at 95°C (t = 3 min), followed by 40 cycles of 95°C (t = 5 s), 60°C (t = 15 s) and 72°C (t = 20 s) and melting curve analysis.

### RT-qPCR data analysis

Raw data were pre-processed with the Real-Time PCR analysis software v4.1.3 (Fluidigm); unspecific values were deleted based on melting-curve analysis. Further processing was done in GenEx v6.0.1 (MultiD Analyses AB): Cq value cutoff of 28 was applied; gDNA background was substracted using ValidPrime^164^ (Tataa Biocenter); data were normalized to the mean expression of 5 reference genes (*Actb, Gapdh, Ppia, Ywhaz, Tubb5*); outliers were deleted (within group Grubbs test, p < 0.05) and a gene was considered undetected for given group if more than 75% values per group were missing; technical replicates (RT and FACS) were averaged; if appropriate, missing data were inputed on a within-group basis and remaining missing data were replaced with Cq_max_ +2 for tissue samples or Cq_max_+0.5 for sorted cells. Of note, the RT-qPCR and RNA-Seq data showed high correlation (Pearson r • 0.942; Figure S13).

Temporal expression of individual genes was first analyzed with two-way ANOVA in R project v3.6.0 using time-point and age as predictor variables, then differences between time-points were tested separately for each age by one-way ANOVA and post-hoc t-tests using *emmeans* package v1.3.5. P-values were adjusted with Benjamini-Hochberg method. Temporal expression of groups of cellular marker genes was first analyzed using linear mixed model in R project v3.6.0 (*lmerTest* package v3.1^154^) with random gene intercept using two-factor design (time-point including control, age) with interaction, then differences between time-points were tested separately for each age. Significance was tested by Satterthwaite’s method. Post-hoc pairwise t-tests were performed using *emmeans* package v1.3.5 and p-values were adjusted by Holm method. Temporal expression of IFN-I pathway was analyzed in similar steps using random slope and random intercept mixed model with 23 interferon-stimulated genes as response variables. IFN-I pathway expression in sorted cells was analyzed in the same way with two-factor design (age, injury) with interaction, using only detected ISGs per each cell type. Differential expression of individual genes (relative to age-matched control) in sorted cells was tested in GenEx v6.0.1 (MultiD Analyses AB) using ANOVA with Bonferroni’s post-hoc test for selected pairwise comparisons and p-values were corrected using Benjamini-Hochberg method (Table S2).

## Acknowledgements

This study was supported by: Czech Science Foundation (16-10214S and 19-02046S); Institutional support (RVO 86652036 and BIOCEV CZ.1.05/1.1.00/02.0109).

## Author contributions

P.An. prepared sequencing libraries, analyzed and interpreted the data, wrote the manuscript and prepared all figures. D.K., J.T. and J.K. performed animal surgeries, prepared cell suspensions and performed FACS experiments. D.Z. and E.R. performed RNA isolation and RT-qPCR experiments. P.Ab. performed raw read pre-processing, mapping and DESeq2 analysis. D.K. M.A. M.K. and L.V. commented on the manuscript. M.K. M.A. and L.V. conceived and supervised the study. All authors reviewed the manuscript.

## Supplementary material

Supplementary material includes Figures S1-S13; Tables S1 and S2; Supplementary excel files containing gene expression matrix, DESeq2 results, gene membership in described gene sets and sets collected from the literature, and searchable excel file for visualization of user-defined genes.

## Availability of data and materials

Raw data will be deposited to public repository and will be made available upon publication of the article. Processed data are available as supplementary files.

