## Supplementary material for "Decoding the transcriptional response to ischemic stroke in young and aged mouse brain": Figures S1 - S13 and Table S1

### Contents:

Figure S1
Figure S2
Figure S3
Figure S4
Figure S5
Figure S6
Figure S7
Figure S8
Figure S9
Figure S10
Figure S11
Figure S12
Figure S13
Table S1

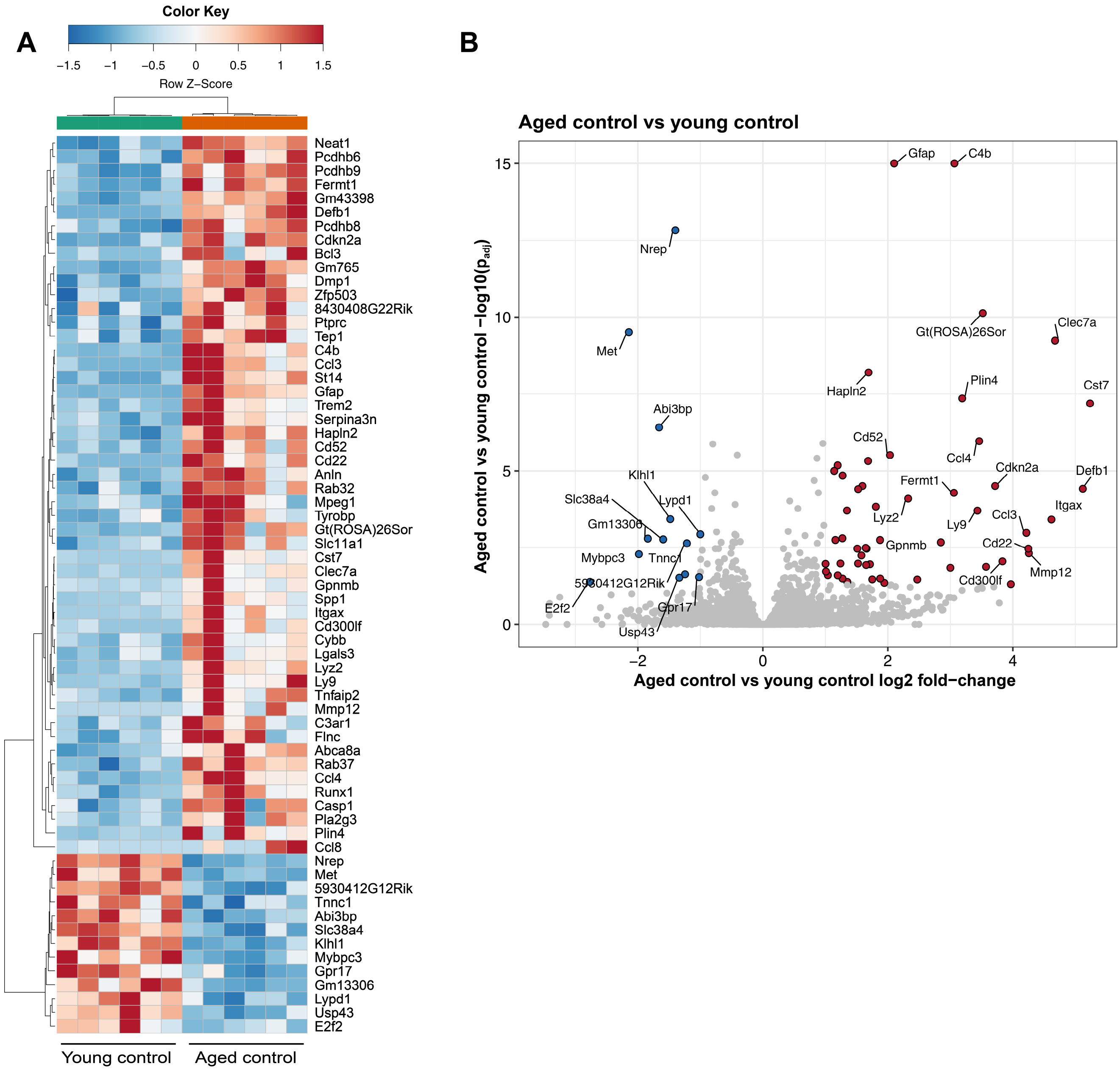


**Figure S1. Differential expression during normal aging**

A) Heatmap showing significantly differentially expressed genes between aged (18 months) and young (3 months) control mice ($|\log_2 FC|>1$; $p_{adj}<0.05$).

B) Volcano plot comparing differential expression between aged and young control mice.

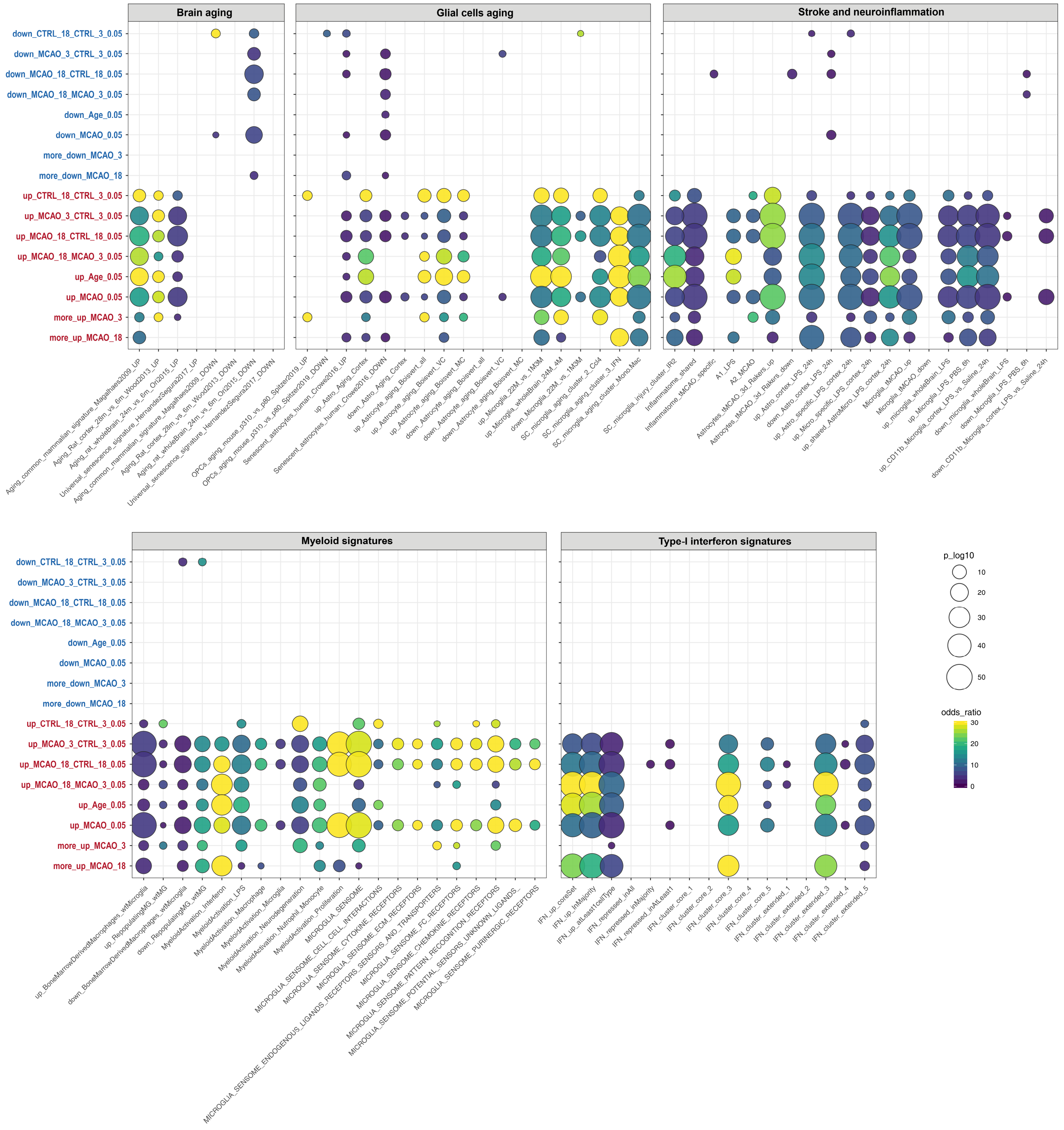


**Figure S2. Overlap of sets of differentially expressed genes with transcriptional signatures collected from literature**
See Table S1 for gene-set descriptions.

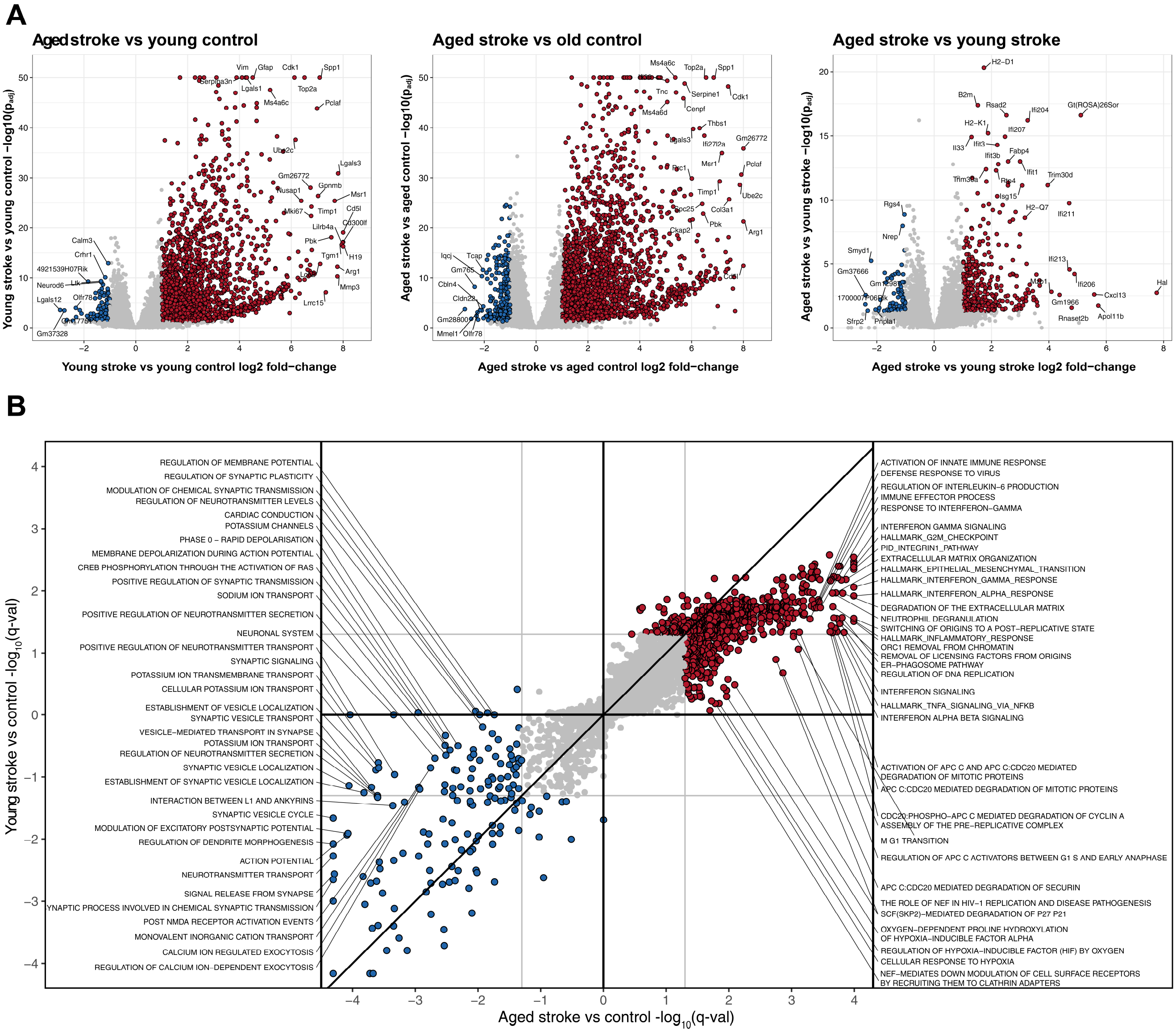

**Figure S3. Comparison of differentially expressed genes and gene sets after stroke between young and aged mice**
A) Volcano plots showing differential gene expression for selected pairwise comparisons.
B) Scatter plot comparing stroke-induced alteration of gene sets from several pathway databases (including Gene Ontology, MsigDB, Panther, Reactome, NetPath, HumanCyc) in young and aged mice. Pathways wih q-val < 0.05 are highlighted in color. Sign depicts UP (+) or DOWN (-) regulation. See also Figure S5.

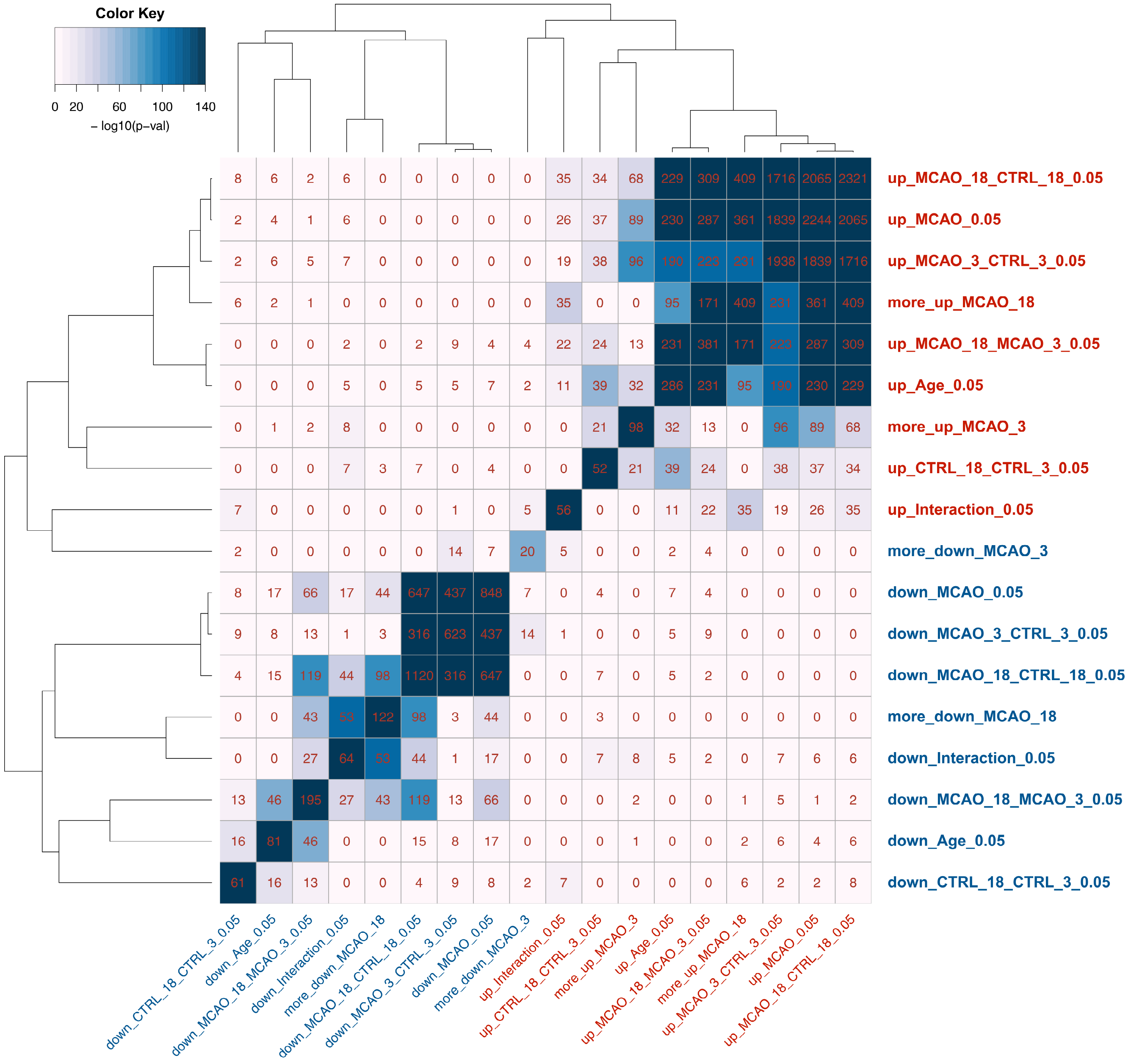


**Figure S4. Clustered heatmap showing overlap between pairs of sets of differentially expressed genes.** Number of intersecting genes is shown in the heatmap.

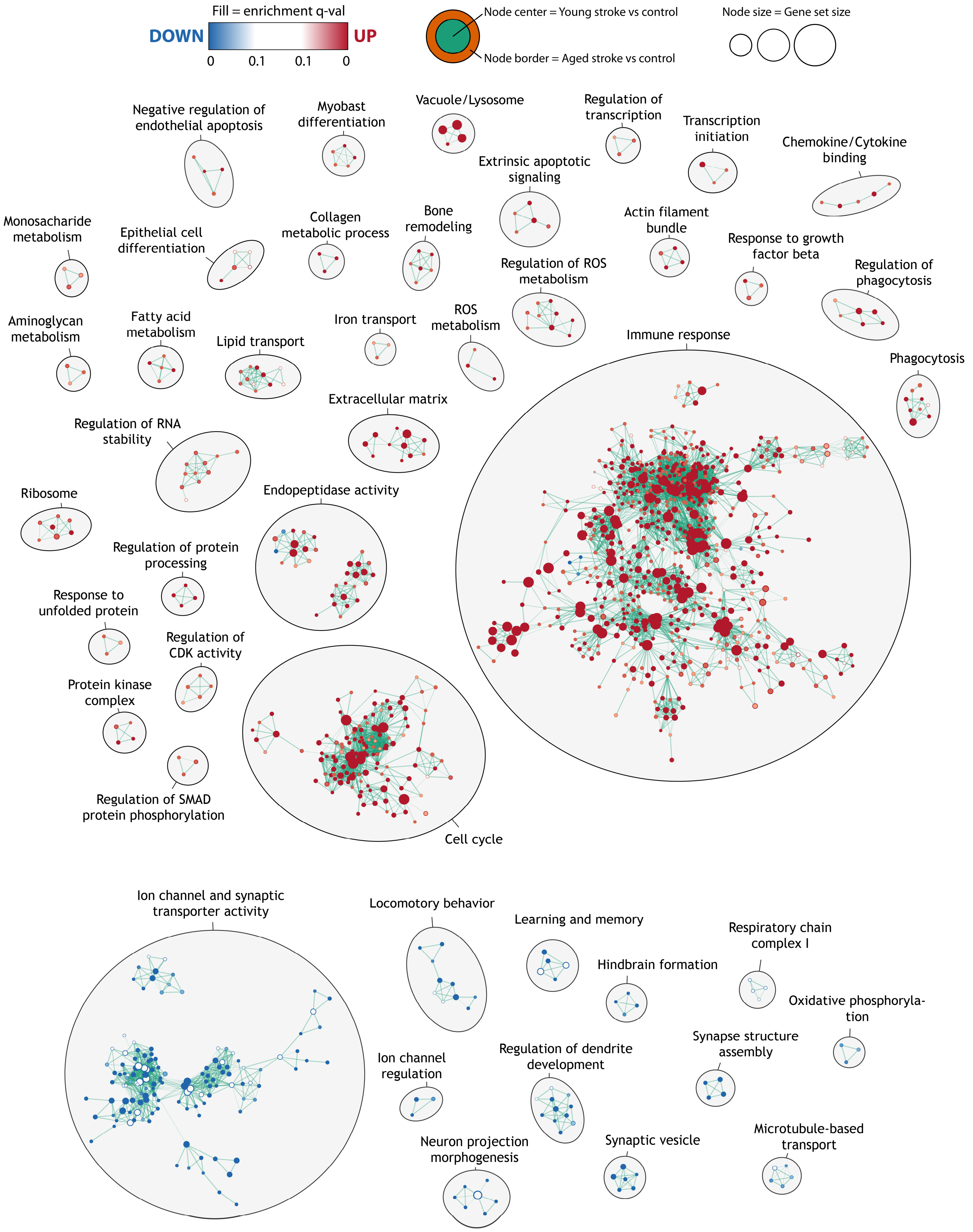


**Figure S5. Enrichment map of significantly UP- or DOWN-regulated gene ontology terms after stroke in young and aged mice.** Nodes represent gene sets. Highly similar gene sets are connected by edges, grouped in sub-clusters and annotated manually.

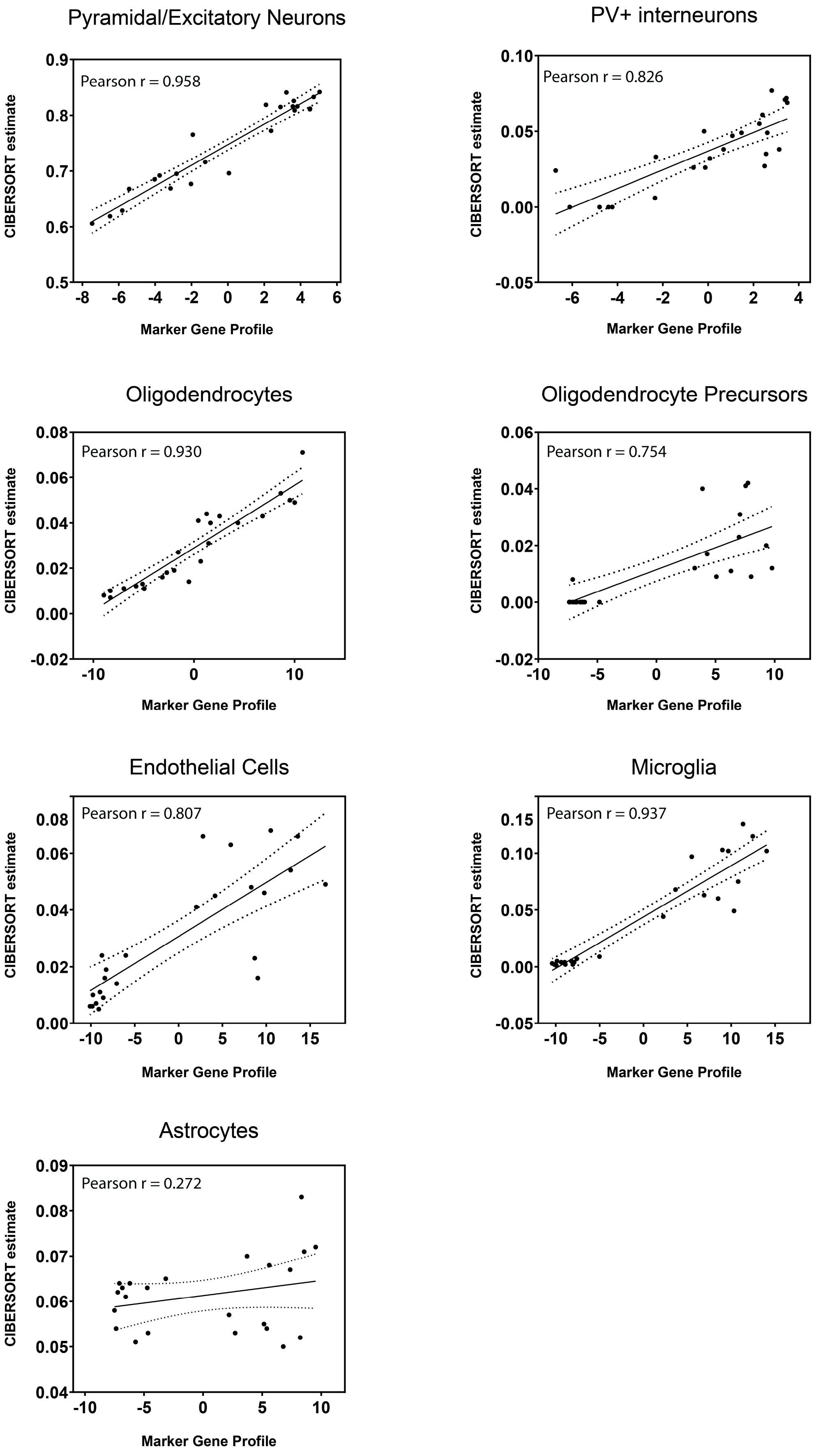


**Figure S6. Concordance between MGP-based and CIBERSORT-based estimates of relative cell proportions.** X-axis shows the loading scores of the first principal component of marker-gene expression determined by *markerGeneProfile* R package (Mancarci et al., 2017). Y-axis shows the relative cell proportions according to CIBERSORT analysis (Newman et al., 2015).

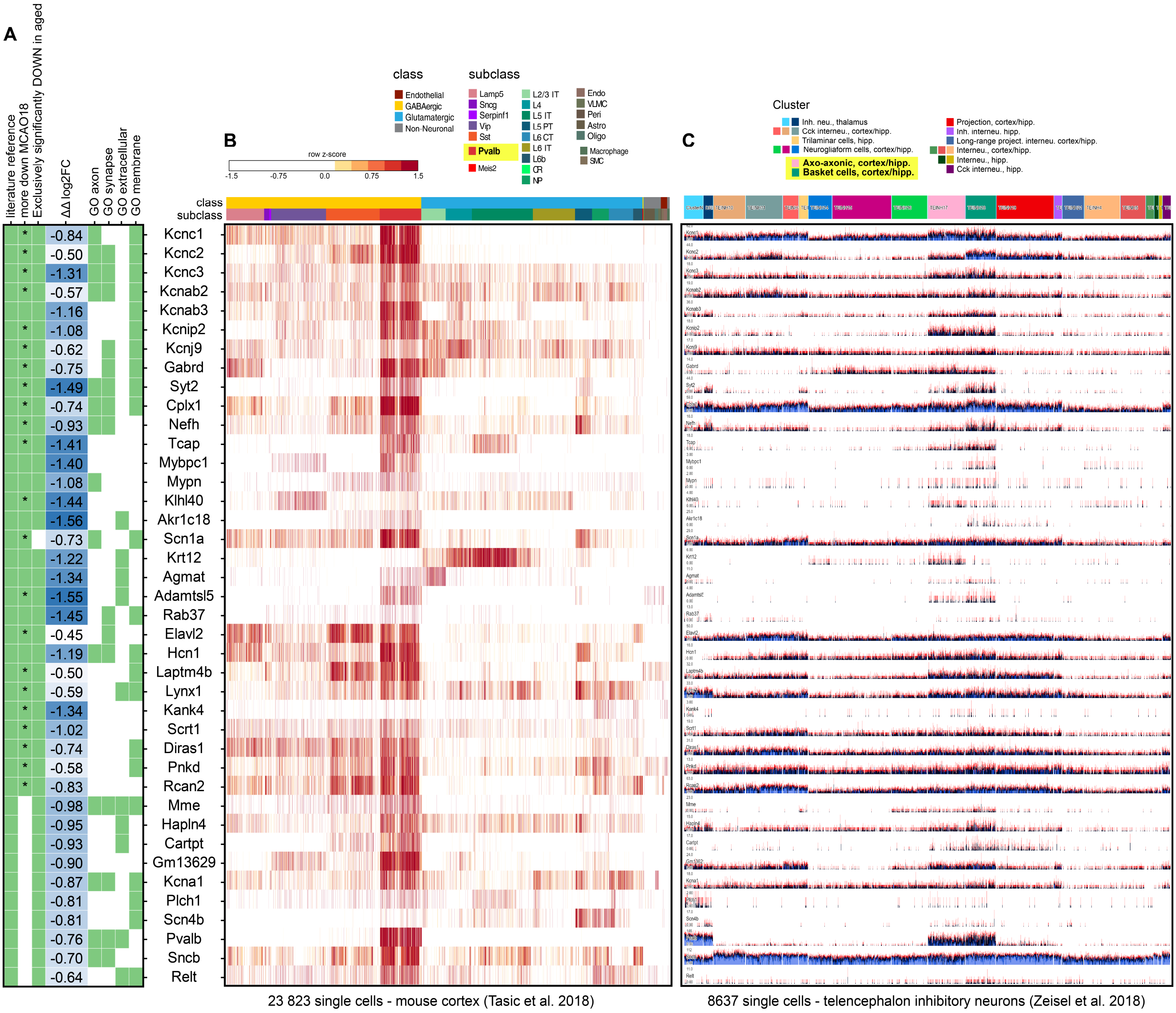


**Figure S7. Genes with greater downregulation after stroke in aged mice („more down MCAO18“) are enriched in PV+ GABAergic interneurons**

A) At least 30 out of 122 genes from “more up MCAO18” gene set as well as other genes significantly downregulated by stroke only in aged mice link to PV+ interneurons through direct literature references. Asterisk depicts significant aging-stroke interaction (DESeq2, p-adj <0.1). Binarized mapping to gene ontology terms “axon”, “synapse”, “membrane” and “extracellular” is also shown.

B) Heatmap showing gene expression in mouse cortex single-cell dataset from Tasic et al., 2018.

C) Sparkline barplot showing single-cell gene expression in telencephalon interneurons from Zeisel et al., 2018.

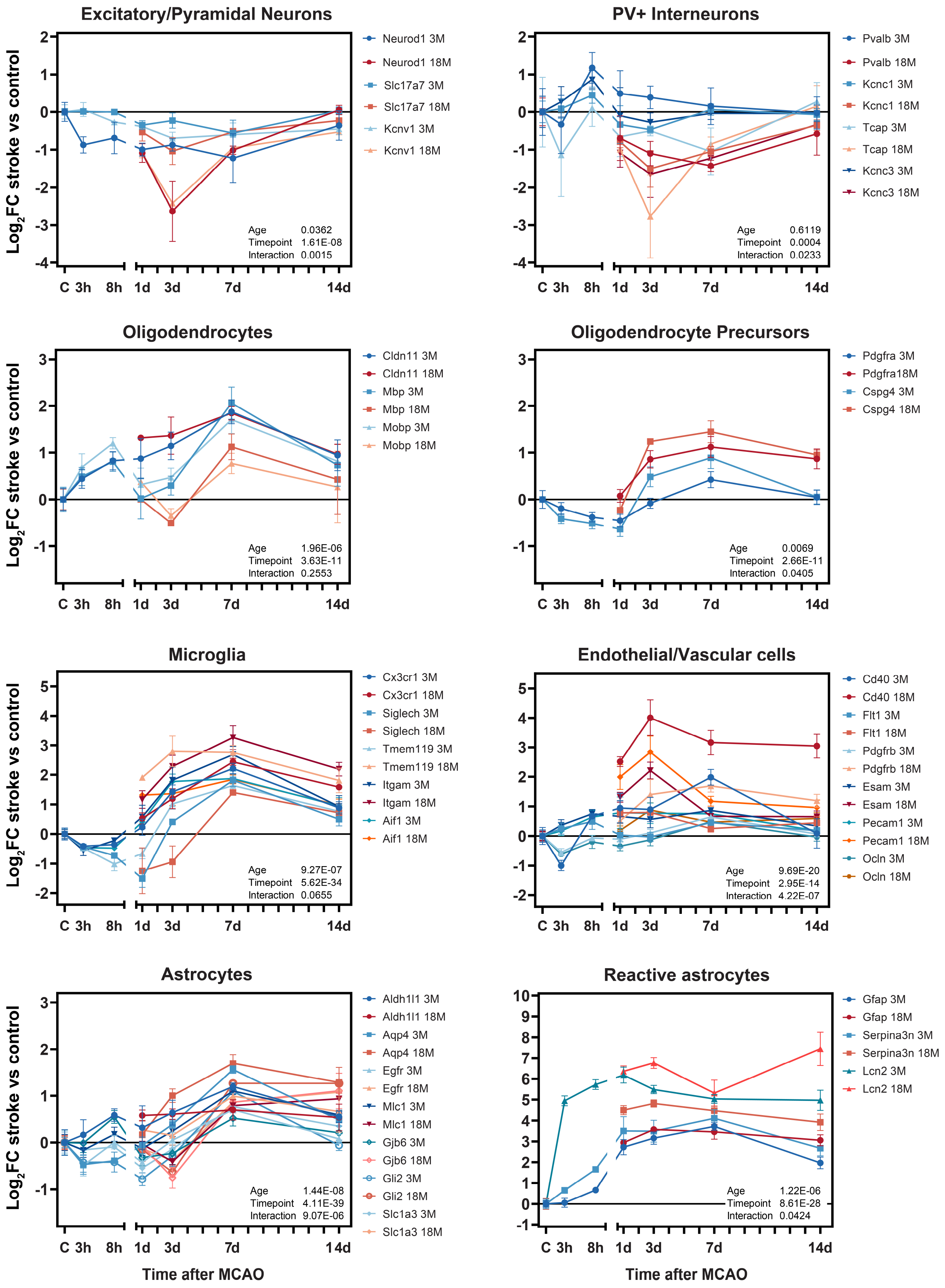


**Figure S8. Temporal expression of selected cell type marker genes after stroke in young and aged mice determined by RT-qPCR.** Young mice shown in shades of blue, aged mice in shades of red. Note the break in the x-axis. P-values for each factor and their interaction are shown at the bottom right corner (linear mixed model).

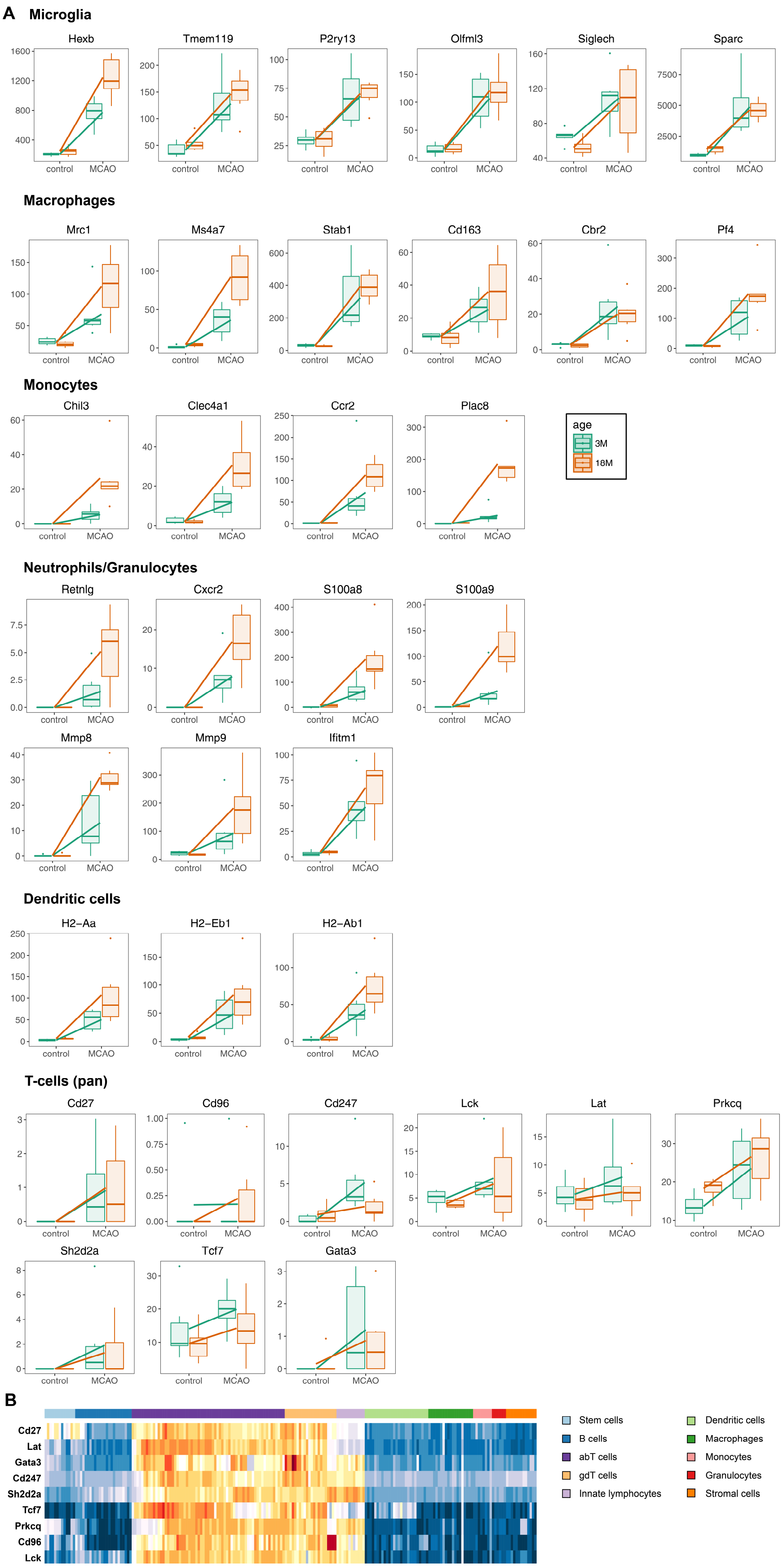


**Figure S9. Expression of selected marker genes of major leukocyte populations 3 days after stroke**

A) Expression boxplots of individual marker genes selected from ImmGen database (www.immgen.org). Lines connect group means. Values on y-axis are in normalized counts.

B) Heatmap from ImmGen database showing expression of selected T-cell pan-marker genes between various leukocyte populations.

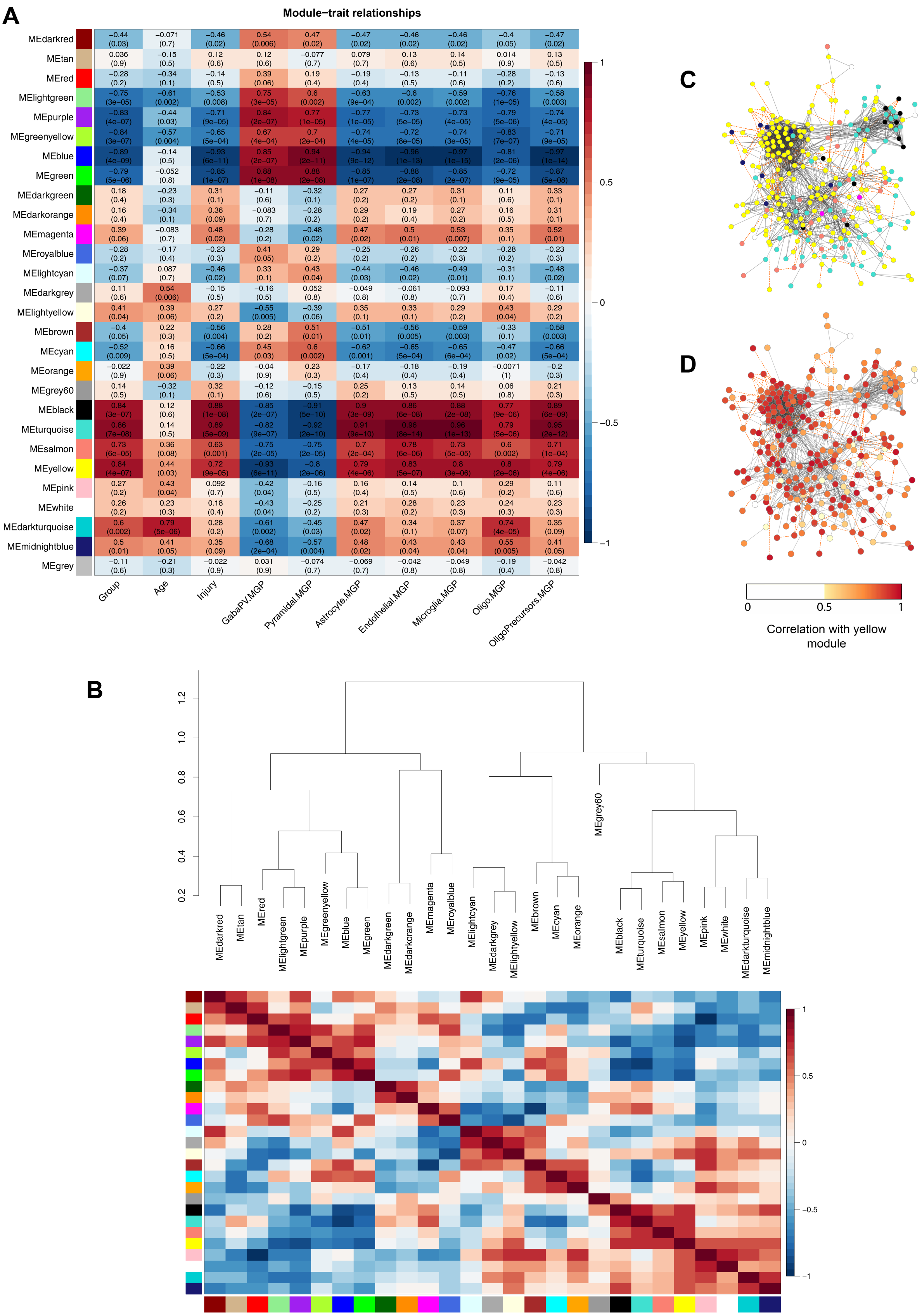


### Figure S10. WGCNA and protein-interaction network characteristics

A) Heatmap showing correlation between WGCNA module eigengenes and deconvoluted cell-specific profiles as well as with age and injury status. Pearson correlation coefficients with p-values are shown.
B) Clustered heatmap showing correlation of WGCNA module eigengenes.
C) Protein interaction network constructed from “more up MCAO18” gene set. Genes are colored according to WGCNA module membership.
D) Same as (C), but genes are colored according to their Pearson correlation with yellow WGCNA module´s eigengene.

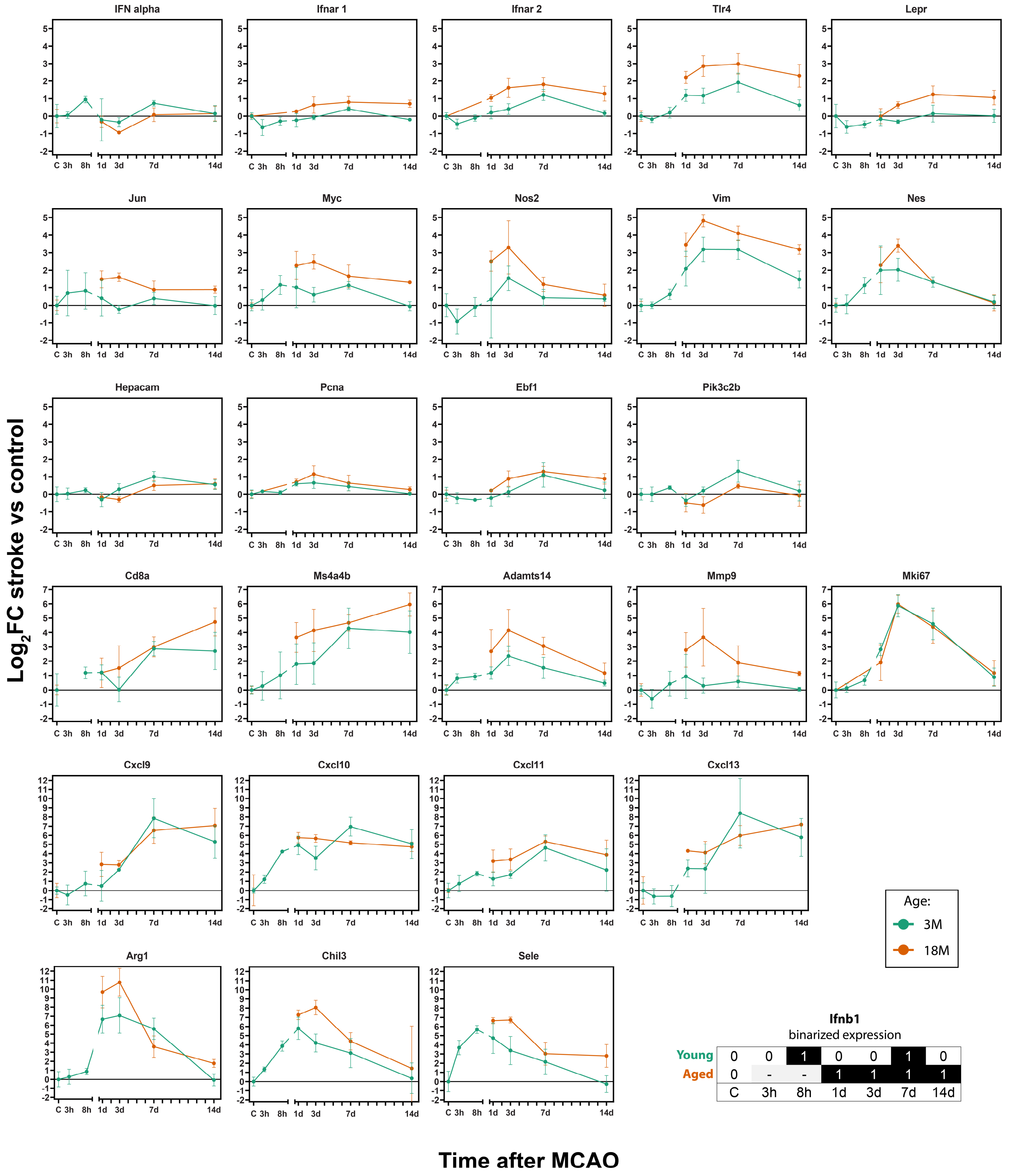


| | | | | | | | |
|---|---|---|---|---|---|---|---|
| Young | 0 | 0 | 1 | 0 | 0 | 1 | 0 |
| Aged | 0 | - | - | 1 | 1 | 1 | 1 |
| | C | 3h | 8h | 1d | 3d | 7d | 14d |

**Figure S11. Temporal expression of selected genes after stroke in young and aged mice.** Note the break in the x-axis. Expression of IFN alpha was measured with degenerate primers targeting the majority of known IFN alpha genes. Expression of *Ifnb1* is binarized as detected (1) or not detected (0) due to Cq values at the limit of detection (bottom right). See Table S2 for results of statistical testing.

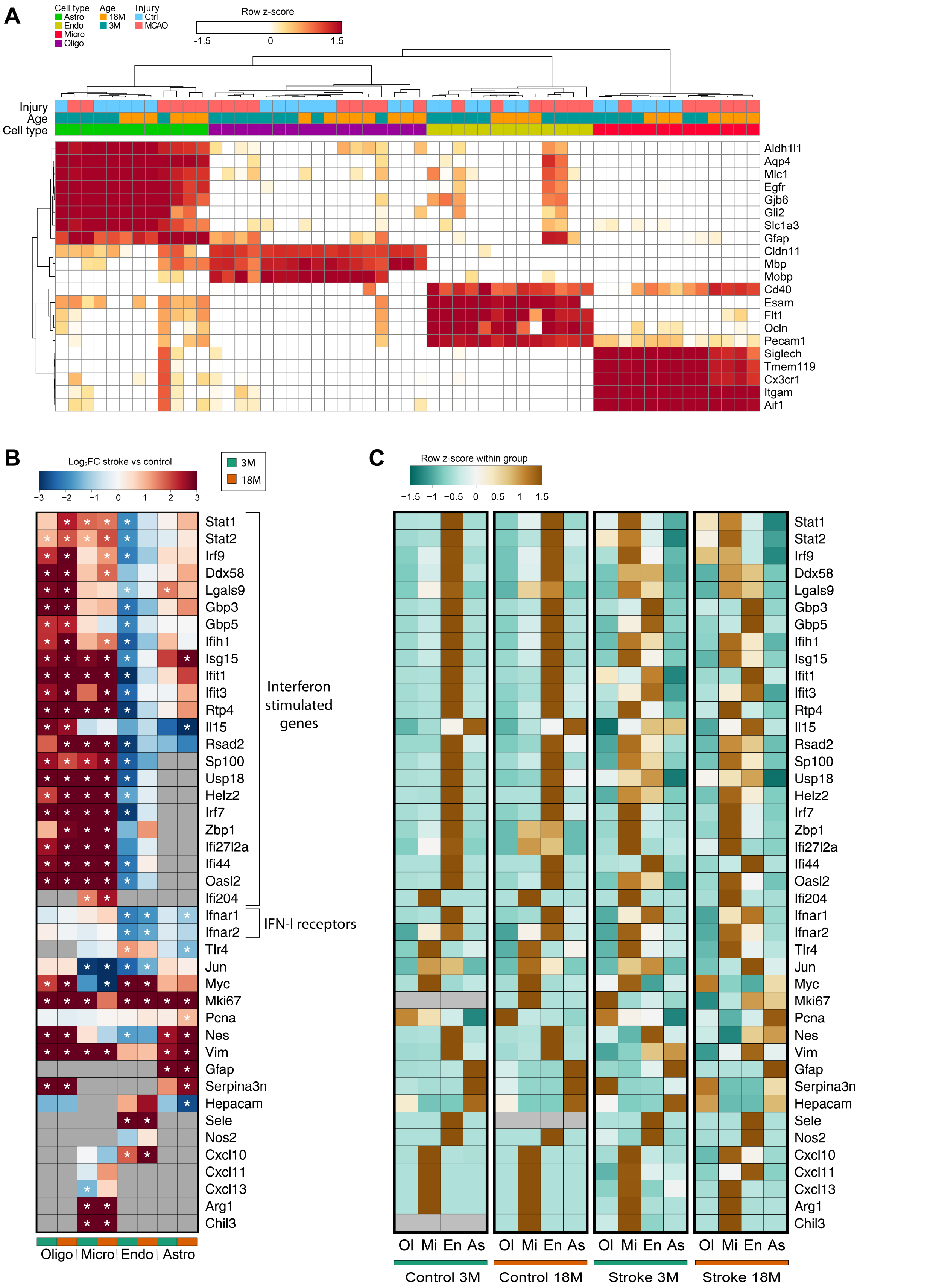


**Figure S12. Cell-specific expression analysis of selected genes after stroke in young and aged mice**

A) Heatmap showing relative expression of cell-type marker genes between individual samples.

B) Heatmap showing log2 fold-change$_{\text{stroke vs control}}$ for each age group and each cell-type separately. Asterisks show significant difference (stroke vs control; p-adj < 0.05; ANOVA with Bonferroni post-hoc test). Signals not detected are shown in grey.

C) Heatmap showing relative expression between cell-types, separately for each group.

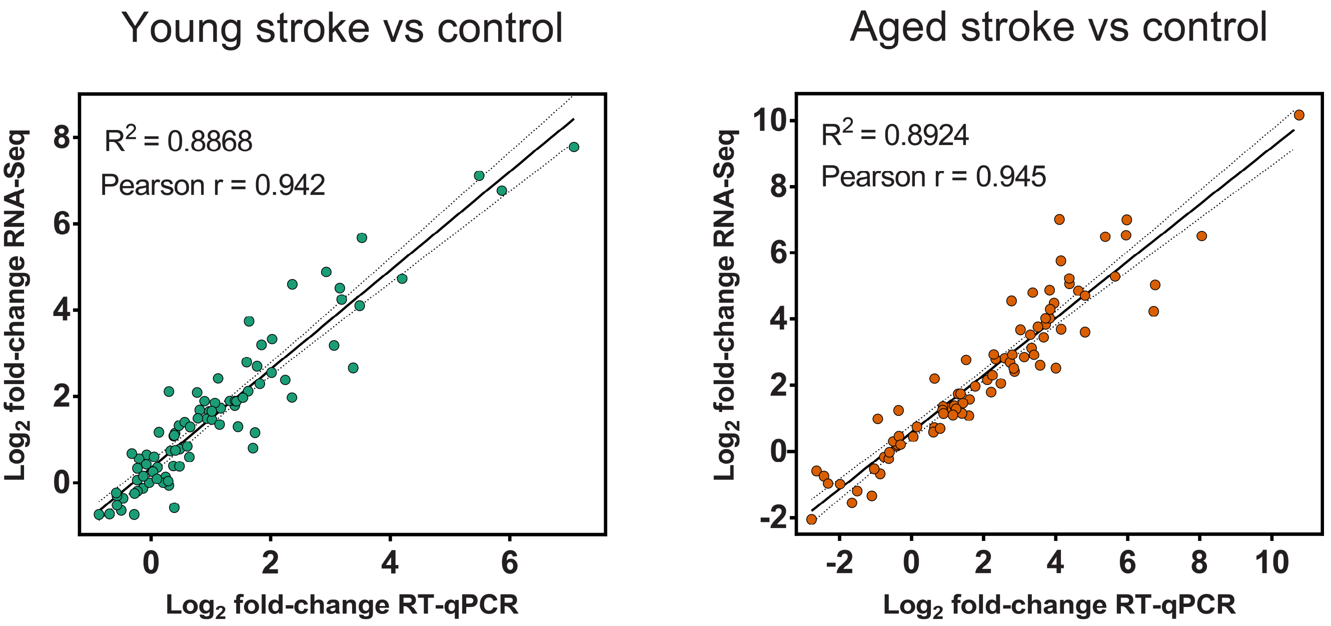


**Figure S13. Concordance between RNA-Seq and RT-qPCR**

Pearson correlation coefficient indicates the concordance of the $log_2FC$ at 3 days after stroke measured by two technologies using two independent sets of samples.

### Table S1. Legend for the gene sets collected from the literature and the gene sets described in this study

| Gene set name | Description | Reference |
|---|---|---|
| Mancarci_cell_markers: | Mouse cortex cell markers from Mancarci et al. 2017 | (Mancarci et al., 2017) |
| McKenzie_cell_markers_.BRETIGEA. | Brain cell markers from McKenzie et al. 2018 | (McKenzie et al., 2018) |
| Barres_Cell_Types | Brain cell markers from Zhang et al., as reanalyzed by Friedman et al. 2018 | (Friedman et al., 2018; Zhang et al., 2014) |
| ABA_cell_types: | Brain cell markers from Allen Brain Atlas, originally published by Tasic et al. 2016, as reanalyzed by Friedman et al. 2018 | (Friedman et al., 2018; Tasic et al., 2016) |
| Han_cell_type: | Brain cell markers from Han et al. 2018 | (Han et al., 2018) |
| SC_cell_clusters_Tasic_2018 | Marker genes of CNS cell clusters („subclass“) according to Tasic et al. 2018. For Glutamatergic cells, subclasses are lumped together (using the higher hierarchy „class“), therefore pan-glutamatergic genes are reported. | (Tasic et al., 2018) |
| SC_pan_cell_clusters_Tasic_2018 | Pan markers of GABAergic vs all other cells and pan-GLUTamatergic vs all other cells according to Tasic et al. 2018 | (Tasic et al., 2018) |
| GABAergic_scRNAseq_markers_Paul_2017 | Markers of GABAergic interneuron subtypes according to Paul et al. 2017 | (Paul et al., 2017) |
| ImmGen_immune_markers: | Cell-enriched genes of immune cell types, according to ImmGen project | www.immgen.org |
| Khoury_Sensome | Microglial sensome, as defined by Hickman et al. 2013 Sensome refers to distinct transcriptional signature of microglia, a unique cluster of transcripts encoding proteins for sensing endogenous ligands and microbes. | (Hickman et al., 2013) |
| Barres_RAmarker | Markers of reactive astrocytes (vs homeostatic astrocytes), either A1/LPS specific (A1_LPS). MCAO/A2 specific (A2_MCAO) or pan-reactive (A_reactive), as defined by Zamanian et al. 2017 and Liddelow et al. 2012 | (Liddelow et al., 2017; Zamanian et al., 2012) |
| Astrocytes_tMCAO_3d_Rakers | genes up or down-regulated in astrocytes after tMCAO (un-adjusted pvalue < 0.05; absolute log2FC > 1) as published by Rakers et al. 2018 | (Rakers et al., 2018) |
| Extracellular.Matrix | Genes that map to Gene Ontology-Cellular Component term „extracellular matrix“ | http://geneontology.org/ |
| Wang_inflammatome | Genes that are part of inflammation transcriptional signature common to 11 disease models (shared) or specific only for tMCAO model (tMCAO_specific) as defined by Wang et al. 2012 (note, that both, upregulated and downregulated genes are part of this list, since original dataset did not contain the info on the sign of regulation). | (Wang et al., 2012) |
| Myeloid_Activation_Fine | Stable co-expressed gene modules found by meta-analysis of transcriptional profiles of brain myeloid cells across various disease/aging models by Friedman et al. 2018 Finer resolution of modules. | (Friedman et al., 2018) |
| Myeloid_Activation_Coarse | Stable co-expressed gene modules found by meta-analysis of transcriptional profiles of brain myeloid cells across various disease/aging models by Friedman et al. 2018 Coarse resolution of modules. | (Friedman et al., 2018) |
| Microglia_tMCAO.Sham | Genes up or down regulated in purified microglia after transient MCAO. Originally published by Arumugam et al. 2017 and reanalyzed by Friedman et al. 2018 | (Arumugam et al., 2017; Friedman et al., 2018) |
| Microglia_aging_22M_vs_1M.3M | Genes up or down regulated in purified microglia from mouse cortex of 22 months old animals VS 1 and 3 month old animals. Originally published by Grabert et al. 2016 and reanalyzed by Friedman et al. 2018 | (Friedman et al., 2018; Grabert et al., 2016) |
| Microglia_aging_wholeBrain_24M_4M | Genes up or down regulated in purified microglia from whole brain of 24 months old animals VS 4 month old animals, as published by Holtman et al. 2015 | (Holtman et al., 2015) |
| SC_microglia_aging_cluster_2_Ccl4 | Marker genes of single-cell microglial clusters published by Hammond et al. 2018 – two aging clusters (OA2, OA3,), monocyte/macrophage cluster (mono/mac) and injury-responsive cluster (IR2). Only markers with log2FC > 1 were considered. | (Hammond et al., 2019) |
| SC_microglia_aging_cluster_3_IFN | | |
| SC_microglia_aging_cluster_Mono.Mac | | |
| SC_microglia_injury_cluster_IR2 | | |

| | | |
|---|---|---|
| microglia_sex_specific_enrichment_cortex | Genes differentially expressed (abs. log2FC >1, padj < 0.05) between male and female murine microglia as published by Guneykaya et al. 2018 | (Guneykaya et al., 2018) |
| BoneMarrowDerivedMacrophages_wtMicroglia | Genes up or down regulated in macrophages derived from bone marrow VS wild-type CNS resident microglia as published by Lavin et al. and reanalyzed by Friedman et al. 2018 | (Friedman et al., 2018; Lavin et al., 2014) |
| RepopulatingMG_wtMG | Genes up or down regulated in repopulating microglia after depletion VS wild-type CNS resident microglia as published by Bruttger et al. 2015 and reanalyzed by Friedman et al. 2018 | (Bruttger et al., 2015; Friedman et al., 2018) |
| LPS_cortex_24h_Micro_Astro_specific | Genes up or down regulated in purified microglia/astrocytes 24h after LPS from mouse cortex as published by Srinivasan et al. Astro_specific = genes changed in astrocytes only, Micro_specific = in microglia only. Shared = in both, astrocytes and microglia. | (Srinivasan et al., 2016) |
| Microglia_LPS_PBS_6h | Genes up or down regulated in purified microglia 6h after LPS injection as originally published by Erny et al. 2015 and reanalyzed by Friedman et al. 2018 | (Erny et al., 2015; Friedman et al., 2018) |
| microglia_wholeBrain_LPS | Genes up or down regulated in purified microglia from whole brain of 4 months old animals 4h after LPS injection as published by Holtman et al. 2015 | (Holtman et al., 2015) |
| ISG | ISG(=Interferon Signature Genes) genes up or down regulated after interferon alpha administration as published by Mostafavi et al. 2016 Up_core refers to core set of genes induced in all 11 immune cell types investigated. Up_majority refers to set upregulated in at least 6 out of 11 immune cell types. Up_atLeast1 refers to set of genes upregulated in at least 1 cell type. | (Mostafavi et al., 2016) |
| IFN_cluster_extended | Distinct regulatory clusters of the interferon signalling network, as published by Mostafavi et al. 2016 | (Mostafavi et al., 2016) |
| IFN_regulator | Whether or not a gene is considered a regulator of interferon signalling network as published by Mostafavi et al. 2016 | (Mostafavi et al., 2016) |
| Universal_senescence_signature_Hernandez.Segura2017 | Genes in the universal signature of the senescent cells (up or down regulated relative no normal cells) as determined by meta-analysis by Hernandez-Segura et al. 2017 | (Hernandez-Segura et al., 2017) |
| Aging_common_mammalian_signature_Magalh.es2009 | Genes that are part of common mammalian gene expression signature of aged cells as determined by meta-analysis by de Magalhães et al. 2009 | (de Magalhães et al., 2009) |
| Aging_rat_wholeBrain_24m_vs_6m_Ori2015 | Genes up or down-regulated in rat whole brain isolated from 24 month old vs 6 month old rats (p-adj <0.05; absolute log2FC > 0) as published by Ori et al. 2015. Data from transcriptomic analysis by RNA-Seq were taken. | (Ori et al., 2015) |
| Aging_Rat_cortex_28m_vs_6m_Wood2013 | Genes up or down-regulated in rat cerebral cortex isolated from 28 month-old rats vs 12 month-old rats (padj < 0.05; absolute log2FC > 0.85) as published by Wood et al. 2013 | (Wood et al., 2013) |
| OPCs.Pdgfra.._aging_mouse_p310_vs_p80_Spitzer2019 | Genes up or down-regulated in murine oligodendrocyte precursor cells isolated from p310 vs p80 mice as published by Spitzer et al. 2019 | (Spitzer et al., 2019) |
| Senescent_astrocytes_human_Crowe2016 | Genes up or down-regulated (padj < 0.05; absolute log2FC >2) in human fetal astrocytes after senescence induction by oxidative stress as published by Crowe et al. 2016 | (Crowe et al., 2016) |
| Astro_Aging_Cortex | Genes up or down regulated in mouse cortex of 2 year old mice vs 10 week old mice as published by Clarke et al. 2018 | (Clarke et al., 2018) |
| Astro_Aging_Hippocampus | Genes up or down regulated in mouse hippocampus of 2 year old mice vs 10 week old mice as published by Clarke et al. 2018 | (Clarke et al., 2018) |
| Astro_Aging_Striatum | Genes up or down regulated in mouse hippocampus of 2 year old mice vs 10 week old mice as published by Clarke et al. 2018 | (Clarke et al., 2018) |
| Astrocyte_aging_Boisvert_all | Genes up or down regulated in mouse motor cortex, visual cortex, hypothalamus and cerebellum of 2 year old mice vs 4 month old mice as published by Boisvert et al. 2018 | (Boisvert et al., 2018) |
| Astrocyte_aging_Boisvert_CB | Genes up or down regulated in mouse cerebellum of 2 year old mice vs 4 month old mice as published by Boisvert et al. 2018 | (Boisvert et al., 2018) |
| Astrocyte_aging_Boisvert_HTH | Genes up or down regulated in mouse hypothalamus of 2 year old mice vs 4 month old mice as published by Boisvert et al. 2018 | (Boisvert et al., 2018) |
| Astrocyte_aging_Boisvert_MC | Genes up or down regulated in mouse motor cortex of 2 year old mice vs 4 month old mice as published by Boisvert et al. 2018 | (Boisvert et al., 2018) |
| Astrocyte_aging_Boisvert_VC | Genes up or down regulated in mouse visual cortex of 2 year old mice vs 4 month old mice as published by Boisvert et al. 2018 | (Boisvert et al., 2018) |

| | | |
|---|---|---|
| ModuleMembership | Membership of a gene in co-expression modules from WGCNA analysis | this study |
| down_CTRL_18_CTRL_3_0.05 | Genes downregulated in aged controls vs young controls (p-adj < 0.05; log2FC < 0) | this study |
| down_MCAO_3_CTRL_3_0.05 | Genes downregulated in young strokes vs young controls (p-adj < 0.05; log2FC < -0.65) | this study |
| down_MCAO_18_CTRL_18_0.05 | Genes downregulated in aged strokes vs aged controls (p-adj < 0.05; log2FC < -0.65) | this study |
| down_MCAO_18_MCAO_3_0.05 | Genes downregulated in aged strokes vs young strokes (p-adj < 0.05; log2FC < -0.65) | this study |
| down_Age_0.05 | Genes downregulated with aging (considering two-factor DESeq2 analysis; p-adj < 0.05; log2FC < -0.65) | this study |
| down_MCAO_0.05 | Genes downregulated with stroke (considering two-factor DESeq2 analysis; p-adj < 0.05; log2FC < -0.65) | this study |
| down_Interaction_0.05 | Genes with negative $\Delta\Delta$log2FC (aged-young) and Age:MCAO interaction p-adj < 0.05 | this study |
| more_down_MCAO3 | Genes downregulated in young strokes vs young controls (p-adj < 0.01; log2FC < -0.65) AND having significant positive interaction ($\Delta\Delta$log2FC > 0 (aged$_{stroke\ vs\ control}$-young$_{stroke\ vs\ control}$); Age:MCAO interaction p-adj < 0.1) AND/OR having $\Delta\Delta$log2FC > 1 (aged$_{stroke\ vs\ control}$-young$_{stroke\ vs\ control}$) | this study |
| more_down_MCAO18 | Genes downregulated in aged strokes vs aged controls (p-adj < 0.01; log2FC < -0.65) AND having significant negative interaction ($\Delta\Delta$log2FC < 0 (aged$_{stroke\ vs\ control}$-young$_{stroke\ vs\ control}$); Age:MCAO interaction p-adj < 0.1) AND/OR having $\Delta\Delta$log2FC < -1 (aged$_{stroke\ vs\ control}$-young$_{stroke\ vs\ control}$) | this study |
| up_CTRL_18_CTRL_3_0.05 | Genes upregulated in aged controls vs young controls (p-adj < 0.05; log2FC > 1) | this study |
| up_MCAO_3_CTRL_3_0.05 | Genes upregulated in young strokes vs young controls (p-adj < 0.05; log2FC > 1) | this study |
| up_MCAO_18_CTRL_18_0.05 | Genes upregulated in aged strokes vs aged controls (p-adj < 0.05; log2FC > 1) | this study |
| up_MCAO_18_MCAO_3_0.05 | Genes upregulated in aged strokes vs young strokes (p-adj < 0.05; log2FC > 1) | this study |
| up_Age_0.05 | Genes upregulated with aging (considering two-factor DESeq2 analysis; p-adj < 0.05; log2FC > 1) | this study |
| up_MCAO_0.05 | Genes upregulated with stroke (considering two-factor DESeq2 analysis; p-adj < 0.05; log2FC > 1) | this study |
| up_Interaction_0.05 | Genes with positive $\Delta\Delta$log2FC (aged$_{stroke\ vs\ control}$-young$_{stroke\ vs\ control}$) AND Age:MCAO interaction p-adj < 0.05 | this study |
| more_up_MCAO3 | Genes upregulated in young strokes vs young controls (p-adj < 0.01; log2FC > 1) AND having significant negative interaction ($\Delta\Delta$log2FC < 0 (aged$_{stroke\ vs\ control}$-young$_{stroke\ vs\ control}$); Age:MCAO interaction p-adj < 0.1) AND/OR having $\Delta\Delta$log2FC < -1 (aged$_{stroke\ vs\ control}$-young$_{stroke\ vs\ control}$) | this study |
| more_up_MCAO18 | Genes upregulated in aged strokes vs aged controls (p-adj < 0.01; log2FC > 1) AND having significant positive interaction ($\Delta\Delta$log2FC > 0 (aged$_{stroke\ vs\ control}$-young$_{stroke\ vs\ control}$); Age:MCAO interaction p-adj < 0.1) AND/OR having $\Delta\Delta$log2FC > 1 (aged$_{stroke\ vs\ control}$-young$_{stroke\ vs\ control}$) | this study |
